# Retraction ATPase motors from three orthologous type IVa pilus systems support promiscuous retraction of the *Vibrio cholerae* competence pilus

**DOI:** 10.1101/2021.10.23.465551

**Authors:** Evan Couser, Jennifer L. Chlebek, Ankur B. Dalia

## Abstract

Bacterial surface appendages called type IVa pili (T4aP) promote diverse activities including DNA uptake, twitching motility, and virulence. These activities rely on the ability of T4aP to dynamically extend and retract from the cell surface. Dynamic extension relies on a motor ATPase commonly called PilB. Most T4aP also rely on specific motor ATPases, commonly called PilT and PilU, to dynamically and forcefully retract. Here, we systematically assess whether motor ATPases from three orthologous T4aP can functionally complement *Vibrio cholerae* mutants that lack their endogenous motors. We found that the PilT and PilU retraction ATPases from the three T4aP systems tested are promiscuous and promote retraction of the *V. cholerae* competence T4aP despite a high degree of sequence divergence. In contrast, the orthologous extension ATPases from the same T4aP systems were not able to mediate extension of the *V. cholerae* competence T4aP despite exhibiting a similar degree of sequence divergence. Also, we show that one of the PilT orthologs characterized does not support PilU-dependent retraction and provide some data to indicate that the C-terminus of PilT is important for PilU-dependent retraction. Together, our data suggest that retraction ATPases may have maintained a high degree of promiscuity for promoting retraction of T4aP, while extension ATPases may have evolved to become specific for their cognate systems.

**IMPORTANCE:** One way that bacteria interact with their environments is via hair-like appendages called type IVa pili (T4aP). These appendages dynamically extend and retract from the cell surface via the action of distinct ATPase motors. T4aP are present in diverse bacterial species. Here, we demonstrate that retraction motors from three T4aP are promiscuous, and capable of promoting retraction of a heterologous T4aP system. By contrast, the extension ATPase motors from these same T4aP systems are specific and cannot promote extension of a heterologous T4aP. Thus, these results suggest that T4aP extension may be more tightly regulated compared to T4aP retraction.

## INTRODUCTION

Type IVa pilus systems (T4aP) are encoded by diverse bacterial species. These systems promote the dynamic extension and retraction of filamentous appendages called pili from the bacterial surface. Through this dynamic activity, T4aP carry out a variety of functions including twitching motility, surface attachment, biofilm formation, virulence, and DNA uptake (1). Dynamic extension of all T4aP relies on the activity of a dedicated motor ATPase, which is commonly called PilB (2). The gene that encodes this motor is generally found in close proximity to other T4aP structural genes. Some systems also have an additional accessory extension ATPase motor called TfpB (3). Dynamic retraction of T4aP canonically relies on a dedicated retraction ATPase called PilT. Additionally, many T4aP systems contain an accessory retraction ATPase called PilU. PilU is a PilT-dependent retraction ATPase that is required for forceful retraction of the competence T4aP in *V. cholerae* (4) and is required for twitching motility in other bacterial species (5). Unlike the extension motor, the genes encoding retraction motors are generally not located near other T4aP structural genes. These hexameric extension and retraction motors are thought to function via a mechanical interaction with the pilus platform, which together take pilin subunits from the inner membrane and polymerize them into a filament during pilus extension or depolymerize these subunits and return them to the inner membrane during pilus retraction. Conformational changes in the ATPase motors due to ATP binding and hydrolysis are thought to power these processes (6). A previous study demonstrated that the PilT retraction motor from *Pseudomonas aeruginosa* was able to complement a *V. cholerae pilT* mutant (7). However, a systematic analysis to assess motor promiscuity among T4aP is lacking.

*Vibrio cholerae* is a facultative pathogen that encodes a competence T4aP, which promotes DNA uptake for natural transformation (8, 9). *Acinetobacter baylyi* and *Pseudomonas stutzeri* encode functionally analogous competence T4aP that also promote horizontal gene transfer (3, 10–12). All of these T4aP systems encode canonical PilB, PilT, and PilU motor ATPases. For this reason, *A. baylyi* and *P. stutzeri* were selected for this study. The T4aP of *A. baylyi* also encodes an additional extension motor called TfpB that is required for initiating pilus extension (3). Some strains of *Escherichia coli* also encode an ortholog of PilT (also referred to as *yggR*), though these strains are not predicted to encode a PilU ortholog.

Here, we sought to formally test promiscuity among T4aP extension and retraction ATPases by using the motors from the systems discussed above. We and others have demonstrated that the competence pilus of *Vibrio cholerae* is a highly genetically tractable system for studying T4aP dynamic activity (4, 7, 8, 13). Thus, we assessed promiscuity among T4aP motors by determining if ectopic expression of heterologous motors could functionally complement *V. cholerae* mutants that lack their endogenous motor ATPases. Our data suggests that while the retraction ATPases PilT and PilU may be promiscuous in their ability to mediate retraction of orthologous T4aP systems, extension ATPases display a high level of specificity for their cognate T4aP. Furthermore, our data identify a PilT ortholog that supports ATPase-dependent retraction, but not PilU-dependent retraction.

## RESULTS

### Retraction ATPase orthologs are promiscuous among T4aP systems, while extension ATPase orthologs are not

Because previous studies have shown that retraction ATPases from T4aP systems may be promiscuous (7), we first wanted to determine if this characteristic is conserved among other bacterial species. To test this, we ectopically expressed *pilT* from *A. baylyi (pilT_Ab_), P. stutzeri* (*pilT_Ps_*), and *E. coli* (*pilT_Ec_*) in *V. cholerae* strains where the native *pilT* or *pilTU* genes were inactivated (i.e., Δ*pilT* and Δ*pilTU* backgrounds). As a control, we also ectopically expressed *V. cholerae pilT* (*pilT_Vc_*). Competence pili in *V. cholerae* promote DNA uptake during natural transformation, a function that requires both pilus extension and retraction (8, 14). Thus, we functionally assessed competence pilus dynamic activity by performing natural transformation assays. Because PilU is an accessory retraction ATPase motor (4, 7), ectopic expression of *pilT_Vc_* alone is able to rescue the transformation defect of the Δ*pilT* and Δ*pilTU* strains (**Fig. 1A**). Furthermore, we observed that ectopic expression of *pilT_Ab_*, *pilT_Ps_*, and *pilT_Ec_*, all fully complemented natural transformation of Δ*pilT* and Δ*pilTU V. cholerae* (**Fig. 1A**). Western blotting of functional N-terminally FLAG-tagged alleles of the PilT orthologs (4) confirmed that their expression was equal to or greater than native PilT in *V. cholerae* (**Fig. S1**).These data indicate that all of these retraction ATPases are capable of mediating retraction of the *V. cholerae* competence T4aP, suggesting that the promiscuity of PilT retraction ATPases is conserved across these species.

**Fig. 1.**
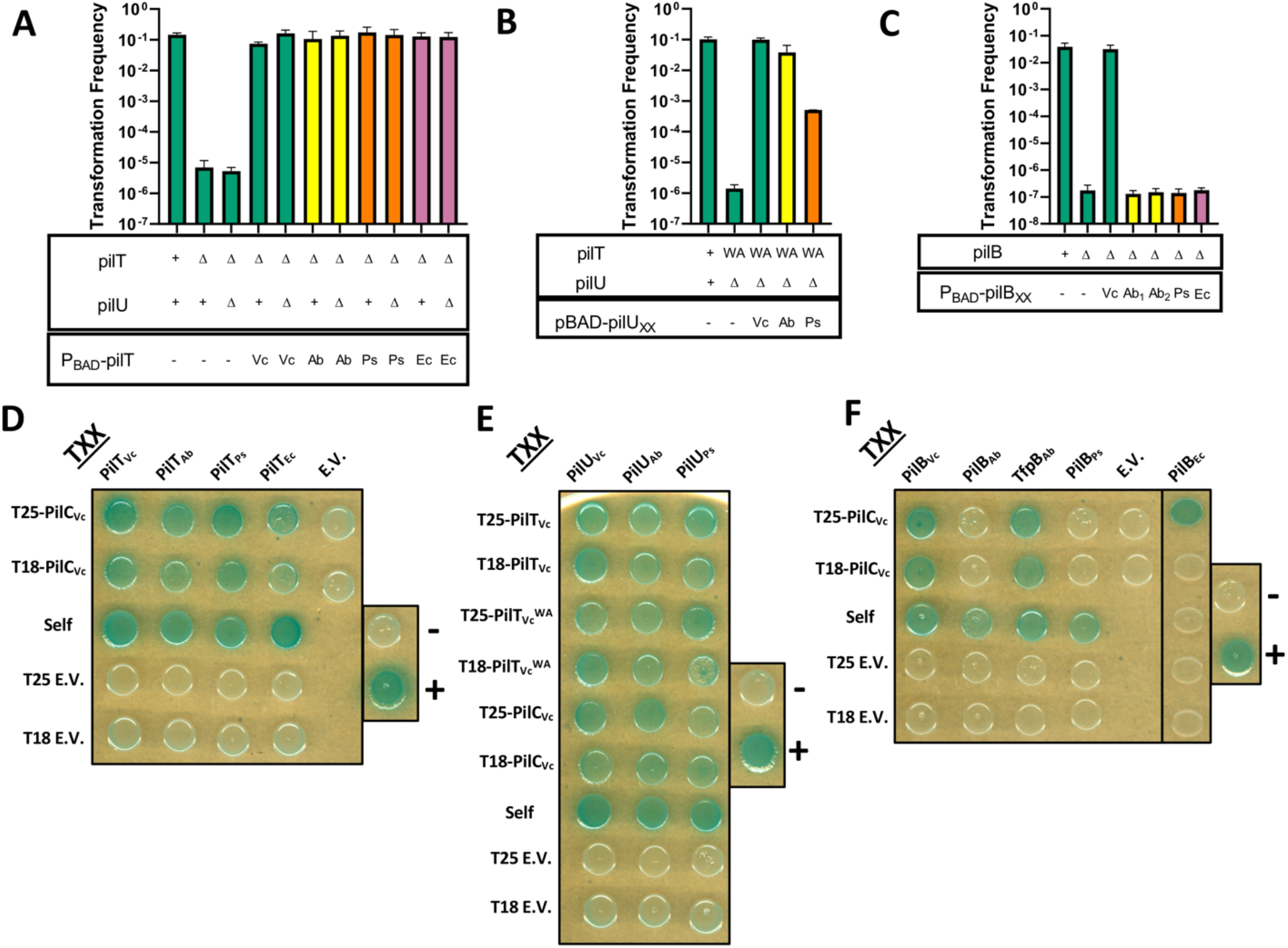
Retraction ATPase orthologs are promiscuous among T4aP systems, while extension ATPase orthologs are not. Natural transformation assays were performed to test whether (**A**) PilT, (**B**) PilU, or (**C**) PilB/TfpB orthologs from 3 heterologous T4aP systems could complement *V. cholerae* mutants lacking their endogenous motor ATPases. The relevant genotypes of strains are indicated below each bar, which includes the status of the native *V. cholerae pilT/pilU/pilB* genes (top) as well as the ectopically expressed motor ATPase allele (induced via P_BAD_) (bottom). The color of the bar corresponds to the species from which the ectopically expressed *motor ATPase* was derived (green bars = *Vc motors,* yellow bars = *Ab* motors, orange bars = *Ps* motors, and purple bars = *Ec* motors). Data are from at least 3 independent biological replicates and shown as the mean ± SD. The limit of detection for transformation frequency assays is ~10^−7^. In **C**, Ab_1_ denotes *pilB_Ab_* and Ab_2_ denotes *tfpB_Ab_*. In **C**, for all samples where the transformation frequency is ~10^−7^, at least 1 replicate contained zero transformants and the limit of detection for that sample is displayed. BACTH assays to test whether the indicated (**D**) PilT, (**E**) PilU, or (**F**) PilB/TfpB alleles can interact with themselves (“self”) or the indicated T4P components (PilC or PilT). For BACTH assays pilus components had an N-terminal fusion to the T25 or T18 fragment of adenylate cyclase as indicated. “^WA^” denotes a Walker A mutation in the tested allele. “+” and “–” denote positive and negative controls for the BACTH assay. “E.V.” denotes an empty vector. Data are representative of at least 2 independent experiments.

In addition to the dominant retraction ATPase PilT, many T4aP also encode an accessory retraction ATPase called PilU that can promote pilus retraction in a PilT-dependent manner (4, 7). So next, we wanted to test whether PilU from *A. baylyi* and *P. stutzeri* could complement a *V. cholerae pilU* mutant (the *E. coli* strain does not have a *pilU* ortholog). To test this, we ectopically expressed *pilU_Ab_* and *pilU_Ps_* in a *V. cholerae* strain where native *pilT* contained a Walker A mutation (*pilT^K136A^* = *pilT^WA^*) and the native *pilU* gene was inactivated (i.e., Δ*pilU*). Mutation of the invariantly conserved lysine in the Walker A domain (K136A) inhibits the ATPase activity of *pilT* (15), thus, efficient pilus retraction in this background (and correspondingly a high frequency of natural transformation) is reliant on PilU ATPase activity (4, 7). As expected, ectopic expression of *pilU_Vc_* recovered the transformation defect of the *pilT^WA^ ΔpilU* strain to parental levels (**Fig. 1B**). The expression of *pilUAb* also showed almost full recovery of transformation when compared to the parent strain (**Fig. 1B**). Interestingly, ectopic expression of *pilU_Ps_* only partially complemented the transformation defect of this strain (**Fig. 1B**). Western blotting of functional N-terminally FLAG-tagged alleles of the PilU orthologs (4) demonstrated that their expression was equal to or greater than native PilU in *V. cholerae* (**Fig. S1**). This suggested that PilU_Ps_ was either (1) generally a less functional PilU motor, or perhaps (2) specifically less active at promoting pilus retraction in conjunction with PilT_Vc_. To test this further, we ectopically expressed *pilU_Ps_* in a background that also ectopically expressed a Walker A mutant of its cognate PilT (*pilT_Ps_^WA^*). We found that in the presence of PilT_Ps_^WA^, PilU_Ps_ was able to completely rescue natural transformation (**Fig. S2**). Together, this suggests that while PilU is also promiscuous between these T4aP systems, it is perhaps less promiscuous than PilT.

Because we found that retraction ATPases are, indeed, promiscuous between these T4aP, we next wanted to test whether the extension ATPases from these same organisms were similarly promiscuous. To test this, we ectopically expressed *pilB* orthologs from *A. baylyi* (*pilB_Ab_* and *tfpB_Ab_*), *P. stutzeri* (*pilB_Ps_*), and *E. coli* (*pilB_Ec_*; also referred to as *hofB*) in a *V. cholerae ΔpilB* mutant and assessed natural transformation. While *pilB_Vc_* recovered natural transformation as expected, ectopic expression of *pilB_Ab_, tfpB_Ab_, pilB_Ps_,* and *pilB_Ec_* failed to complement the Δ*pilB* mutant (**Fig. 1C**). Importantly, Western blotting of N-terminally FLAG-tagged alleles of all PilB orthologs (14) indicated that expression of each was equal to or greater than native PilB in *V. cholerae* (**Fig. S1**). This suggests that extension ATPase orthologs from these species are not able to mediate extension of the *V. cholerae* competence pilus. These data indicate that there may be a high degree of specificity among extension ATPases, which is in stark contrast to the observed promiscuity among the retraction ATPases studied here.

Extension and retraction ATPases likely interact with the pilus machinery platform protein PilC to promote dynamic pilus activity (16). Thus, the observed promiscuity and/or specificity among these motor ATPases may be attributed to their ability and/or inability to interact with PilC. To test this, we assessed protein-protein interactions genetically using Bacterial Adenylate Cyclase Two-Hybrid (BACTH) assays (17). First, looking at PilT, we found that each of the PilT orthologs could self-interact, which is consistent with the hexameric conformation that likely all pilus motors adopt (**Fig. 1D**). Also, consistent with the observation that all of the PilT orthologs tested support natural transformation, we found that all PilT orthologs could interact with PilC_Vc_ (**Fig. 1D**). Because PilU is a PilT-dependent retraction ATPase, it may need to interact with both PilT and PilC to support pilus retraction. Indeed, all of the PilU orthologs could interact with PilT_Vc_, PilT_Vc_^WA^, and PilC_Vc_, which is consistent with the ability of each PilU ortholog to support some degree of natural transformation (**Fig. 1E**). For the PilB orthologs, we found that PilBvc could interact with PilCvc as expected (**Fig. 1F**). Additionally, each of the PilB orthologs, apart from PilB_Ec_, could self-interact, which is consistent with the hexameric conformation that these proteins likely adopt (**Fig. 1F**). Interestingly, Pi1B_Ab_, TfpB_Ab_, PilB_Ps_, and PilB_Ec_ varied in their ability to interact with PilC_Vc_. PilB_Ab_, PilB_Ps_, could not interact with PilCvc and PilB_Ec_ only interacted with PilC_Vc_ in one direction (**Fig. 1F**), which could suggest that these proteins fail to support extension because they cannot interact properly with the platform. However, TfpB_Ab_ could interact with PilC_Vc_ (**Fig. 1F**), but still failed to support natural transformation (**Fig. 1C**). These data indicate that while interactions between PilB and PilC may be required for extension of the competence pilus, these interactions are not sufficient to promote extension.

### The PilT ortholog from E. coli cannot support PilU-mediated retraction

Above, our results indicate that PilT orthologs from distinct T4aP can promote retraction of the *V. cholerae* competence pilus in the absence of native PilT_Vc_ and PilU_Vc_. PilT, however, can also support PilU-dependent retraction, in a manner that is not dependent on PilT ATPase activity (4). So next, we wanted to test if the PilT orthologs among the studied T4aP could support PilU-dependent retraction in conjunction with PilU_Vc_. To test this, Walker A mutants of each *PilT* gene (*pilT_Vc_^WA^, pilT_Ab_^WA^, pilT_Ps_^WA^, pilT_Ec_^WA^*) were ectopically expressed in a Δ*pilT* background (i.e., where *pilU_Vc_* is intact) and a Δ*pilTU* background. As expected, the ectopic expression of *PilT_Vc_^WA^* fully restored the transformation efficiency of a Δ*PilT* background (since *pilU_Vc_* is intact) (**Fig. 2A**). Importantly, transformation was not restored by ectopic expression of any Walker A *PilT* alleles in a Δ*PilTU* background (**Fig. 2A**), which indicates that these ATPase defective alleles of PilT only support PilU-dependent retraction and cannot promote retraction on their own. When *pilT_Ab_^WA^* and *pilT_Ps_^WA^* were ectopically expressed in the Δ*pilT* background, transformation was rescued to parental levels (**Fig. 2A**). These data suggest that PilU_Vc_ is able to mediate retraction of the *V. cholerae* competence pilus in a PilT_Ab_- and PilT_Ps_-dependent manner, which further demonstrates the promiscuity among these retraction ATPases. Ectopic expression of the Walker A mutant allele of *pilT_Ec_*, however, did not restore natural transformation in the Δ*PilT* background (**Fig. 2A**). This suggests that PilT_Ec_^WA^ is incapable of promoting PilU-dependent retraction in the presence of PilU_Vc_.

**Fig. 2.**
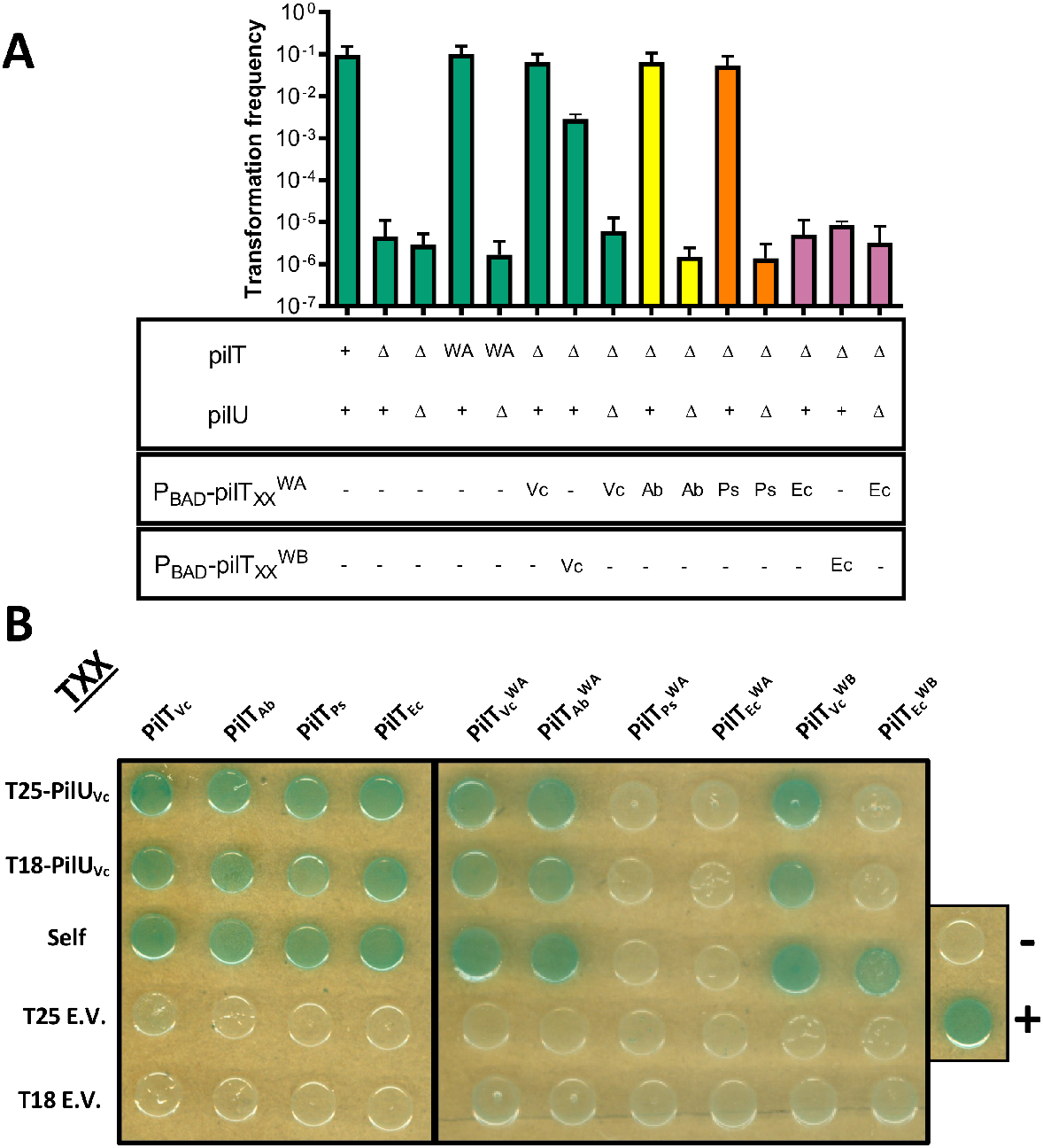
The PilT homolog from E. coli cannot support PilU-mediated retraction. (**A**) Natural transformation assays were performed to test whether retraction ATPase PilT mutants from 3 heterologous T4aP systems could complement *V. cholerae* mutants lacking their endogenous motor ATPases. The relevant genotypes of strains are indicated below each bar, which includes the status of the native *pilT/pilU* genes as well as the ectopically expressed motor ATPase allele (induced via P_BAD_). The color of the bar corresponds to the species from which the ectopically expressed *PilT* was derived (green bars = *Vc pilT,* yellow bars = *Ab pilT,* orange bars = *Ps pilT,* and purple bars = *Ec pilT*). Data are from at least 3 independent biological replicates and shown as the mean ± SD. (**B**) BACTH assays to test whether the indicated PilT alleles can interact with themselves (“self”) or PilU_Vc_. For BACTH assays pilus components had an N-terminal fusion to the T25 or T18 fragment of adenylate cyclase as indicated. “^WA^” denotes a mutation in the Walker A domain of the tested allele, while “^WB^” denotes a mutation in the Walker B domain of the tested allele. “+” and “–” denote positive and negative controls for the BACTH assay. “E.V.” denotes an empty vector. Data are representative of at least 2 independent experiments.

The ability of PilU to interact with PilT is likely necessary for PilU-dependent retraction of the competence pilus. As expected, PilT_Vc_, PilT_Ab_, PilT_Ps_, and PilT_Ec_ could self-interact (**Fig. 2B**) as seen previously (**Fig. 1D**), and each allele could also interact with PilU_Vc_ (**Fig. 2B**). Because PilU_Vc_-mediated retraction was possible when *pilT_Vc_^WA^, pilT_Ab_^WA^*, and *pilT_Ps_^WA^* were ectopically expressed, we hypothesized that each of these PilT alleles would be able to self-interact and also interact with PilU. PilT_Vc_^WA^ and PilT_Ab_^WA^ were able to self-interact (**Fig. 2B**), which suggests that the Walker A mutation does not inhibit the ability of these proteins to self-assemble into hexamers. This is consistent with our previous results which demonstrate that purified PilT_Vc_^WA^ can still form hexameric rings (4). Unexpectedly, PilT_Ps_^WA^ and PilT_Ec_^WA^ did not exhibit self-interactions in these BACTH assays (**Fig. 2B**). We also observed that PilU_Vc_ could interact with PilT_Vc_^WA^ and PilT_Ab_^WA^, however, it did not interact with PilT_Ec_^WA^ or PilT_Ps_^WA^ (**Fig. 2B**). The absence of interactions between PilU_Vc_ and PilT_Ec_^WA^ (**Fig. 2B**) is consistent with the inability of PilT_Ec_^WA^ to promote PilU-dependent retraction (**Fig. 2A**). However, the absence of interactions between PilU_Vc_ and PilT_Ps_^WA^ was particularly surprising since this PilT allele can support PilU-dependent retraction when expressed in *V. cholerae* (**Fig. 2A**). Because PilT_Ps_^WA^ supports PilU-dependent retraction in *V. cholerae*, but does not exhibit self-interactions in BACTH assays, this may suggest that self-assembly or hexamerization of PilT is not required for PilU-dependent retraction. However, the inability of PilT_Ps_^WA^ to interact with PilU_Vc_ in these same BACTH assays (**Fig. 2B**), despite the ability of this allele to support PilU-dependent retraction (**Fig. 2A**) makes this difficult to conclude. The observed inability of PilT_Ec_^WA^ and PilT_Ps_^WA^ to self-interact and/or interact with PilU_Vc_ in BACTH assays could either be due to an absence of interactions between these proteins, or could be due to a poor affinity interaction between these proteins that is below the detection limit for the assay.

The molecular mechanism underlying PilU-mediated retraction in a PilT-dependent manner remains unclear. The ability of PilT to self-assemble and form a hexamer may be a prerequisite for PilU-mediated retraction. However, other characteristics of PilT may also be responsible for supporting PilU-mediated retraction. Because *PilT_Ec_^WA^* lacked self-interactions, we constructed an alternative mutant that also disrupted the ATPase activity of PilT_Ec_ by mutating an invariantly conserved glutamate in the Walker B domain of *PilT_Ec_ (PilT_Ec_^E201A^* = *PilT_Ec_^WB^*). Importantly, PilT_Ec_^WB^ exhibited self-interactions in BACTH assays (**Fig. 2B**), suggesting that the Walker B mutation allows for self-assembly and/or a higher affinity between subunits compared to the Walker A mutant. When we ectopically expressed the *PilT_Ec_^WB^* in a Δ*PilT* background (where *pilU* is intact), however, we observed that this mutant did not restore transformation (**Fig. 2A**), which is similar to the *PilT_Ec_^WA^* allele. This indicates that self-assembly of an ATPase defective PilT_Ec_ is not sufficient to promote PilU-dependent retraction. Together, these data suggest that PilT_Ec_ may have unique features that prevents it from promoting PilU-dependent retraction. We hypothesized that analyzing the differences between PilT_Ec_ and the other PilT alleles (PilT_Vc_, PiIT_Ab_, and PilT_Ps_) may help define the features of PilT that allow for PilU-dependent retraction.

### The C-terminus of PilT_Vc_ contributes to PilU-mediated retraction

In order to narrow down the region of PilT_Vc_ that is necessary and/or sufficient to support PilU-dependent retraction, we constructed defined chimeras of *PilT_Vc_* and *PilT_Ec_*. PilT_Vc_ and PilT_Ec_ are 345 residues and 326 residues long, respectively. The chimeras were constructed in intervals of 50 amino acids where the residues of one protein were replaced with the corresponding amino acids of the other. These chimeras were first ectopically expressed in a Δ*PilTU* background to test their ability to mediate retraction of the competence pilus in a manner that is dependent on their ATPase activity – a property that both PilT_Vc_ and PilT_Ec_ shared (**Fig. 1A**). Surprisingly, only three of the chimeras were able to restore parental levels of transformation in the Δ*PilTU* background (**Fig. 3A**). Two of the functional chimeras consisted of portions of the N-terminus of PilT_Vc_ (100 or 200 amino acids) fused to the corresponding C-terminus of PilT_Ec_. The remaining functional chimera contained the first 300 amino acids of PilT_Ec_ fused to the C-terminal 42 amino acids of PilT_Vc_. One potential explanation for this general lack of functionality is that the locations of the junctions between the two proteins were not conducive to generating a functional ATPase motor. PilT is composed of two domains, an N-terminal PAS-like domain and a C-terminal RecA-like ATPase domain which are connected via a short linker (95-102 in PilT_Ec_ and 98-105 in PilT_Vc_). So, the chimeras at 100 residues should be within the linker that connects the two domains of both proteins, and it was particularly surprising that the PilT_Ec_(100) fusion was not functional when the reciprocal PilT_Vc_(100) fusion was functional. The three functional chimeras of PilT obtained are the only alleles that we can be reasonably confident could potentially support PilU-dependent retraction, thus, we only proceeded with characterizing these three chimeras of PilT moving forward. To determine if these three chimeras could support PilU-mediated retraction, we next introduced a Walker A mutation into each allele. None of the *PilT_Vc_* / *PilT_Ec_* chimeras with the Walker A mutation were able to rescue PilU-dependent transformation efficiency in a Δ*PilT* background (**Fig. 3A**). Separately, a Walker B mutation was introduced into these chimeras. Again, none of the 3 chimeras were able to restore PilU-dependent transformation in a Δ*PilT* background (**Fig. 3A**). We reasoned that since PilT_Vc_ is capable of PilU-mediated retraction, that the chimera that is predominantly composed of PilT_Vc_ might help define a portion of the protein that plays an important role in PilU-mediated retraction (i.e. that the part of PilT_Vc_ missing in this chimera plays an important role). PilT_Vc_(200) had the largest portion of PilT_Vc_ and still made a functional retraction motor when ATPase activity was intact (**Fig. 3A**). This chimera had 200 residues of the PilT_Vc_ N-terminus fused to the C-terminal 129 residues of PilT_Ec_. Walker A and Walker B mutants of PilT_Vc_(200) did not support PilU-dependent retraction (**Fig. 3A**). So, we reasoned that the C-terminal 145 residues of PilT_Vc_ (which were replaced with PilT_Ec_ in these chimeras) must be playing an important role in PilU-dependent retraction. However, this does not exclude the possibility that the N-terminus of PilT_Vc_ also plays an important role in promoting PilU-dependent retraction.

**Fig. 3.**
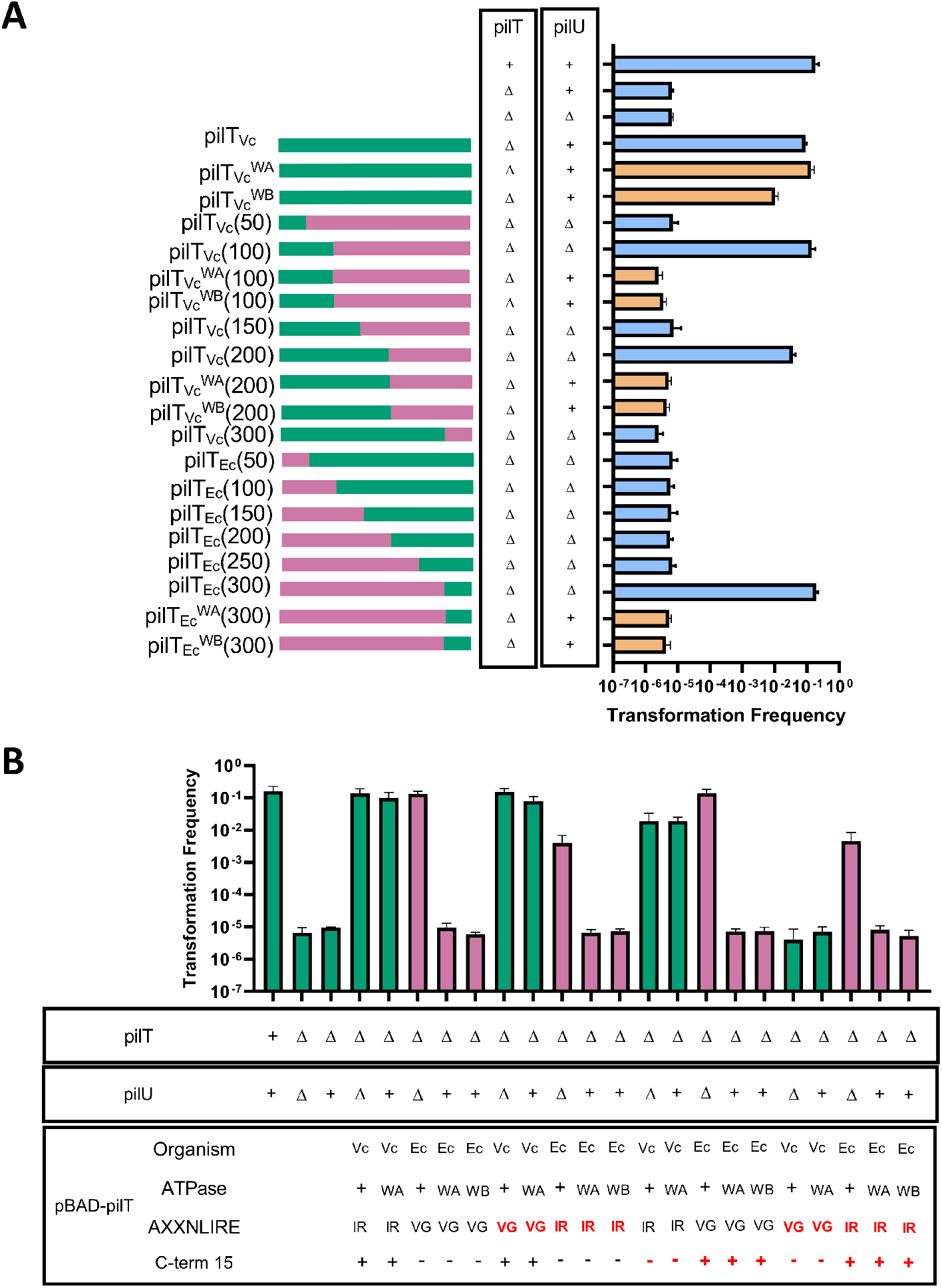
The C-terminus of PilT_Vc_ contributes to PilU-mediated retraction. Natural transformation assays were performed to test whether retraction ATPase chimeras between PilT_Vc_ and PilT_Ec_ can complement *V. cholerae* mutants lacking their endogenous motor ATPases. PilT chimeras either have (**A**) their ATPase activity intact to assess ATPase-dependent retraction, or their ATPase activity inactivated via Walker A or Walker B mutations to assess PilU-mediated retraction. The relevant genotypes of strains are indicated to the left of each bar, which includes the status of the native *pilT/pilU* genes as well as the ectopically expressed motor ATPase chimera (induced via P_BAD_). Diagrams to the left of bars denote the composition of each chimera with the fragments of PilT_Vc_ denoted in green and PilT_Ec_ denoted in pink. For strains that encode an ATPase active *pilT* allele, the bar is blue, while for strains that contain an ATPase inactive *pilT* allele (via a WA or WB mutation as indicated) the bar is tan. The number in parentheses indicates the number of N-terminal amino acids from the PilT allele indicated fused to its PilT ortholog. (**B**) Natural transformation assays were performed to test whether two elements at the C-terminus of PilT_Vc_ (the AIRNLIRE domain and a 15-residue domain at the C-terminus) contribute to PilU-mediated retraction. The relevant genotypes of strains are indicated below each bar, which includes the status of the native *pilT/pilU* genes as well as the ectopically expressed motor ATPase chimera (induced via p_BAD_). Four attributes of each ectopically expressed allele are indicated: the organism from which the PilT allele is derived, the ATPase status of the allele (“+” denotes ATPase intact, “WA” denotes Walker A mutant, and “WB” denotes Walker B mutant), the sequence of the AXXNLIRE domain (IR = AIRNLIRE, VG = AVGNLIRE), and the presence / absence of the C-terminal 15 amino acids. If the attribute changes the natural allele, it is displayed in red text. The color of the bar corresponds to the species from which the ectopically expressed *pilT* was derived (green bars = *Vc pilT* and purple bars = *Ec pilT*). All data are from at least 3 independent biological replicates and shown as the mean ± SD.

In order to further investigate potential components of PilT that are required for supporting PilU-mediated retraction, we constructed an alignment of the 4 PilT orthologs that were studied here. The alignment revealed that the middle section of PilT_Ec_ was conserved relative to all of the other PilT orthologs (**Fig. S3**). However, the N-terminus and the C-terminus of PilT_Ec_ were poorly conserved relative to the other PilT orthologs. Because our data suggests that the C-terminus of PilT_Vc_ may be important for supporting PilU-mediated retraction, that is where we focused our investigation. In particular, we observed that *pilT_Ec_* contains two substitutions in the highly conserved AIRNLIRE domain of PilT (18). To determine if these substitutions were responsible for the inability of PilT_Ec_ to support PilU-mediated retraction (7), we mutated the two residues in *pilT_Ec_* to the residues present in the other PilT orthologs (PilT_Ec_^AvG→AIR^). Conversely, to determine if these changes in PilT_Ec_ contributed to its inability to support PilU-mediated retraction, the residues in *PilT_Vc_* were mutated to the residues present in *PilT_Ec_* (PilT_Vc_^AIRàAvG^). These mutations were introduced into Walker A (WA) or Walker B (WB) mutant alleles of PilT so that we could assess PilU-mediated retraction. The introduction of these mutations into *PilT_Vc_* did not result in the inability of PilT_Vc_ to facilitate PilU-mediated retraction (**Fig. 3B**). Moreover, the restoration of the two AļRNLIRE substitutions in *pilT_Ec_* did not rescue PilU-mediated retraction (**Fig. 3B**). These data indicate that the substitutions in the AIRNLIRE domain of *PilT_Ec_* are not responsible for the inability of pilT_Ec_ to support PilU-mediated retraction.

The alignment of the 4 PilT orthologs also revealed that 15 amino acids in the C-terminal domain of PilT_Ec_ were missing when compared to the other PilT orthologs that did support PilU-mediated retraction (PilT_Vc_, PilT_Ab_, and PilT_Ps_). We hypothesized that the deletion of the last 15 amino acids in PilT_Ec_ could be responsible for the inability of this protein to support PilU-mediated retraction. To test this, we deleted the last 15 amino acids from PilT_Vc_ (*pilT_Vc_*^Δ15aa^)’ Additionally, we fused the last 15 amino acids present at the C-terminus of *pilT_Vc_* to the end of pilT_Ec_ (*PilT_Ec_^+15aa^*). Again, these mutations were generated in Walker A and Walker B mutant alleles of PilT so that we could assess PilU-mediated retraction. The *pilT_Vc_^Δ15aa, WA^* mutant displayed no reduction in transformation (**Fig. 3B**), suggesting that PilT_Vc_^Δ15aa, WA^ is still able to facilitate PilU-mediated retraction. Conversely, the *PilT_Ec_^+15aa, WA^* and *PilT_Ec_^+15aa, WB^* mutations did not restore transformation to parental levels (**Fig. 3B**), which suggests that PilU_Vc_ cannot mediate retraction in the presence of these PilT_Ec_ alleles. Together, these data suggest that the last 15 amino acids present at the C-terminus of PilT_Vc_ is not necessary or sufficient to promote PilU-mediated retraction.

Our results have suggested that, individually, the substitutions present in the conserved AIRNLIRE domain of PilT_Ec_ and the deletion of the last 15 amino acids in the C-terminus are not responsible for the inability of this PilT allele to support PilU-mediated retraction. However, it is possible that the combination of these two factors diminishes PilU-mediated retraction. To test this, we constructed *pilT_Vc_ alleles* that contained both the AIRNLIRE domain substitutions and the C-terminal deletion of 15 amino acids (*piiTv_Vc_*^AIR→AvG,Δ15aa^). Also, we generated a *PilT_Ec_* allele where we restored the AIRNLIRE domain and inserted the terminal 15 amino acids from *PilT_Vc_* onto the C-terminus (*pilT_Ec_*^Avg→AIR, +15aa^). The combination of mutations in *pilT_Vc_* resulted in a loss of function, even in a strain where the ATPase activity of the allele was intact (**Fig. 3B**).

Additionally, the *pilT_Ec_*^AvG→AIR, +15aa, WA^ and *pilT_Ec_*^AvG→AIR, +15aa, WB^ alleles did not restore transformation to parental levels in the Δ*pilT* background, suggesting that PilU was unable to mediate retraction in these backgrounds. This suggests that the combination of the 15 amino acid deletion and the substitutions in the AIRNLIRE domain are not solely responsible for the inability of PilT_Ec_ to support PilU-mediated retraction.

## DISCUSSION

This work demonstrates that the retraction ATPases PilT and PilU are promiscuous among the subset of T4aP systems studied here, and that their functionality extends beyond species boundaries. This contrasts with the fidelity displayed by the extension ATPases from the same T4aP systems. Additionally, our data suggests that the C-terminus of PilT_Vc_ plays a role in supporting PilU-mediated retraction. However, further analysis determined that the genetic differences between PilT_Vc_ and PilT_Ec_ in the AIRNLIRE domain and the extended C-terminus are neither necessary nor sufficient to explain this discrepancy in activity. Because of this, the exact residues in the PilT C-terminus that contribute to PilU-mediated retraction remain elusive. Also, it is likely that the N-terminus of PilT_Vc_ also contributes to PilU-dependent retraction, however, the paucity of functional PilT_Vc_ / PilT_Ec_ chimeras made this difficult to narrow down.

Because the orthologous retraction motor ATPases were overexpressed in this study, we cannot comment on the relative efficiency of the ectopically expressed retraction motors compared to native PilT_Vc_ *and* PilU_Vc_. But it is important to note that similar overexpression of the extension motor orthologs from the same T4aP did not promote dynamic competence pilus activity, which indicates that the overabundance of a motor ATPase is not necessarily sufficient for motor functionality. One possible explanation for the discrepancy in the promiscuity / specificity among motor ATPases could be due to differences in sequence divergence; where retraction motor orthologs are more similar to one another compared to the extension motor orthologs. When we performed this analysis, however, we found that the PilT orthologs ranged from ~62-80% similar (with PilT_Ec_ being the most divergent), the PilU orthologs ranged from ~70-74% similar, and the extension motor orthologs ranged from ~50-66% similar (**Fig. S4**). These data indicate that overall sequence divergence among PilT and PilB orthologs is fairly similar. Thus, suggesting that sequence divergence alone cannot account for the promiscuity / specificity among these motor ATPases.

Why are retraction ATPases promiscuous and extension ATPases specific? One possibility is that it is advantageous for T4aP systems to maintain promiscuous retraction ATPases. T4aP are diverse and retraction ATPases are generally expressed at distinct loci from the other T4aP components. Some T4aP systems are likely transferred between organisms through horizontal gene transfer (19). The inability to retract T4aP is likely disadvantageous in many circumstances since these surface appendages are highly immunogenic and can also be receptors for diverse phages (20). Thus, it is tempting to speculate that one potential advantage to retraction ATPase promiscuity is that this would allow cells that acquire a new T4aP the ability to retract pili from the surface.

Some organisms, like *V. cholerae,* encode multiple T4P - competence pili, mannose sensitive hemagglutinin (MSHA) pili, and toxin co-regulated pili (TCP) - and there are distinct environmental conditions under which each pilus system is likely needed. Competence pili and MSHA pili are likely critical for the survival of *V. cholerae* in its environmental reservoir, while TCP are required for infection of its human host. In fact, it has previously been shown that the inappropriate expression of MSHA pili during the context of an infection attenuates *V. cholerae* (21). A high degree of specificity among the extension ATPase motors for these distinct T4P may represent one mechanism by which cells can exhibit tight control over when these distinct pili are made. In contrast, the retraction of these pili may not require the same level of regulation because pilus extension is so tightly regulated. Indeed, in *V. cholerae,* PilT and PilU promote retraction of both competence pili and MSHA pili (8, 22).

The molecular mechanism underlying the specificity of extension ATPase motors remains unclear. One possibility is that the N-terminal domain of PilB provides specificity. It is thought that the most N-terminal domain of PilB is responsible for interactions with PilM, which is likely required for proper pilus extension (6). An alignment of the extension and retraction motor orthologs showed that the extension motors contain an extended N-terminal domain (~184 residues), called N1D (6) that distinguishes them from the retraction ATPase motors (**Fig. S5**), and we found that much of the sequence divergence among extension ATPases was attributed to the N1D (**Fig. S4**). Thus, it is tempting to speculate that the N1D of extension ATPases plays a critical role in defining the specificity of these motors for their cognate T4aP system.

While our results indicate that retraction motors are promiscuous and extension motors are specific among the T4aP studied here, it remains unclear which of these phenotypes is the exception vs the rule. To distinguish between these possibilities, it would be interesting to test the degree of specificity / promiscuity among other T4aP components. Addressing this question may continue to shed light on the evolutionary pressures that have helped shaped these nearly ubiquitous molecular machines, and will be the focus of future research.

## METHODS

### Bacterial Strains and Growth Conditions

*Vibrio cholerae* strains were grown in LB Miller broth and LB Miller agar plates that contained erythromycin (10 μg/mL), carbenicillin (20 μg/mL), and/or kanamycin (50 μg/mL) when necessary. A list of all strains used in this study can be found in **Table S1**.

### Construction of Mutant Strains and Constructs

All of the *V. cholerae* strains used in this study are derivatives of the El tor isolate E7946. The motor ATPase orthologs from other species were amplified from genomic DNA of the following strains: *Acinetobacter baylyi* ADP1 (23), *Pseudomonas stutzeri* JM300 (24), and *Escherichia coli* MG1655 (25). All mutant constructs were made via splicing-by-overlap extension (SOE) PCR, exactly as previously described (13, 26). A detailed list of all primers used to generate mutant constructs can be found in **Table S2**. Mutant constructs were introduced onto the *V. cholerae* genome by natural transformation and natural cotransformation exactly as previously described (26). Strains were constructed by first inserting the ectopic expression construct of the pilus component into the genome. This was followed by the deletion of the native pilus component. All mutations and ectopic expression constructs were confirmed by PCR and/or sequencing.

### Natural Transformation Assays

In this study, we aimed to address the role of distinct T4aP components on competence pilus function. Thus, we opted to bypass the natural mechanism of competence induction in *V. cholerae* by ectopically expressing the master competence regulator TfoX (*P_tac_-tfoX*) and inactivating the quorum sensing negative regulator LuxO (Δ*luxO*) as previously described (8). Chitin-independent natural transformation assays were performed exactly as previously described (8, 14). Briefly, strains were grown overnight in 3 mL of LB broth containing 100 μM IPTG at 30 °C rolling. Next, cells were subcultured into 3 mL of LB supplemented with 20 mM MgCl_2_, 10 mM CaCl_2_, 100 μM IPTG, and arabinose (when appropriate; 0.05-0.2%) and grown to late-log phase at 30 °C rolling. Next, ~10^8^ cells were transferred to Instant Ocean medium (7 g/L; Aquarium Systems) containing 100 μM IPTG, arabinose (when appropriate; 0.05-0.2%), and 200 ng of a transforming DNA (tDNA) PCR product. Reactions were then incubated statically overnight at 30 °C. The tDNA product used throughout this study targeted a “neutral gene” (the frame-shifted transposase, VC1807) for inactivation (ΔVC1807::Erm^R^) as previously described (27). A negative control, containing no tDNA, was included for each strain in all transformation assays. After the overnight incubation with tDNA, reactions were outgrown by adding 1 mL of LB to each reaction and incubating at 37 °C shaking for 2 hours. After the outgrowth, reactions were plated for quantitative culture to distinguish between transformants (plated on LB + erythromycin) and total viable counts (plated on plain LB). The transformation frequency for each reaction was calculated by dividing the number of transformants by the total viable counts.

### Bacterial Adenylate Cyclase Two-Hybrid Assay

Using PCR, the desired motor ATPase genes were amplified and cloned into the BACTH vectors pKT25 and pUT18C in order to produce N-terminal fusions between the motor ATPase and the T25 and T18 fragments of adenylate cyclase, respectively. Miniprepped plasmids (Qiagen) were then co-transformed into *E. coli* BTH101 cells. As a positive control, T25-zip and T18-zip vectors were cotransformed into BTH101. And as a negative control, the pKT25 and pUT18C empty vectors were cotransformed into BTH101. Transformations were plated onto LB + kanamycin (50 μg/mL) + carbenicillin (100 μg/mL) plates in order to select for transformants that received both plasmids. These transformants were then selected and grown overnight at 30 °C static in LB + kanamycin (50 μg/mL) + carbenicillin (100 μg/mL). Finally, 3 μL of the overnight cultures were spotted onto LB agar plates supplemented with 500 μM IPTG, kanamycin (50 μg/mL), carbenicillin (100 μg/mL), and X-gal (40 μg/mL). These plates were allowed to incubate statically at 30 °C for ~48 hours prior to imaging on a HP Scanjet G4010 flatbed scanner.

### Alignments

The sequences of the genes that encode for the desired proteins were downloaded from GenBank. Sequences were aligned using T-Coffee (28) and the resulting alignments were run through the Sequence Manipulation Suite: Color Align Conservation tool (29) to generate the final figures.

### Pairwise comparisons to assess similarity among ATPase motor orthologs

Pairwise comparisons between ATPase motor orthologs were performed via global alignment using the Needleman-Wunsch algorithm (30). The percent similarities generated by these pairwise comparisons are reported in **Fig. S4**.

### Western Blotting

Strains were grown to late-log phase in LB supplemented with 20 mM MgCl_2_, 10 mM CaCl_2_, 100 μM IPTG, and arabinose (when appropriate) exactly as described above for transformation assays. Cells were washed and concentrated to an OD_600_ = 100 in 0.5X Instant Ocean medium, and then lysed by adding Fastbreak lysis buffer (to 1X), lysozyme (80 μg/mL), and benzonase (25U). Lysed samples were mixed 1:1 with 2X SDS sample buffer [220 mM Tris pH 6.8, 25% glycerol, 1.8% SDS, 0.02% Bromophenol Blue, 5% β-mercaptoethanol] and 3 μL of each sample was then separated on a 15% SDS PAGE gel. The proteins were transferred to a PVDF membrane via electrophoresis, and the membrane was then incubated with the appropriate primary antibody (α-FLAG polyclonal rabbit – for PilT and PilU; or α-FLAG M2 mouse monoclonal – for PilB). A primary incubation with α-RpoA monoclonal mouse (BioLegend) was also completed as a loading control. The blots were then washed and incubated with an appropriate horseradish peroxidase (HRP) conjugated secondary antibody. And blots were developed with Pierce™ ECL Western Blotting Substrate and imaged using a ProteinSimple FluorChem R system.

## ACKNOWLEDGEMENTS

We would like to thank David Kysela for helpful discussions. This work was supported by grant R35GM128674 from the National Institutes of Health (to ABD).

---

**Fig. S1.**
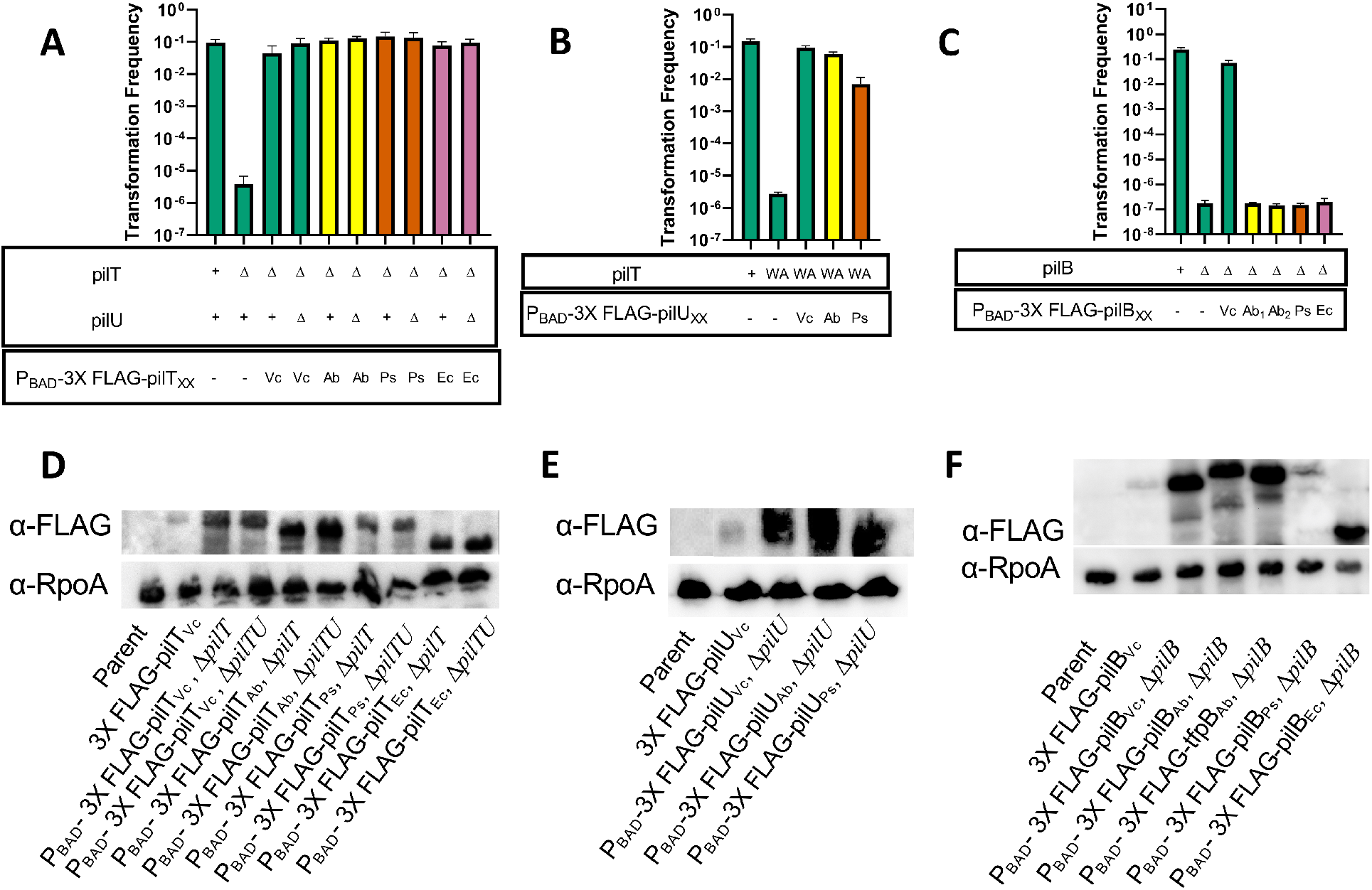
Expression of ectopic constructs is equal to or greater than native levels of expression. Natural transformation assays were performed to confirm the functionality of the N-terminally FLAG-tagged alleles of **(A)** *pilT,* **(B)** *pilU,* and **(C)** *pilB*. The relevant genotypes of strains are indicated below each bar, which includes the status of the native *V. cholerae pilT/pilU/pilB* genes (top) as well as the ectopically expressed motor ATPase allele (induced via P_BAD_) (bottom). The color of the bar corresponds to the species from which the ectopically expressed *motor ATPase* was derived (green bars = *Vc motors,* yellow bars = *Ab* motors, orange bars = *Ps* motors, and purple bars = *Ec* motors). Data are from at least 3 independent biological replicates and shown as the mean ± SD. The limit of detection for transformation frequency assays is ~10^−7^. In **C**, Ab_1_ denotes *pilB_Ab_* and Ab_2_ denotes *tfpB_Ab_*. In **C**, for all samples where the transformation frequency is ~10^−7^, at least 1 replicate contained zero transformants and the limit of detection for that sample is displayed. Western blots to determine the levels of expression of the FLAG-tagged **(D)** *pilT,* **(E)** *pilU,* and **(F)** *pilB* alleles and RpoA as a loading control. The blots show that all ectopically expressed motors were expressed at or above the levels of the native *Vc* motor. Data is representative of at least two independent experiments.

**Fig. S2.**
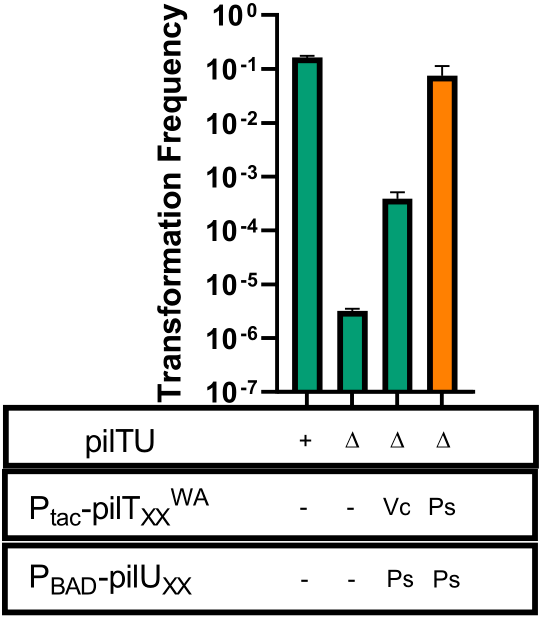
PilU_Ps_ is a fully functional retraction ATPase motor. Natural transformation assays were performed to assess whether PilU_Ps_ can support pilus retraction in conjunction with its cognate PilT_Ps_^WA^ better than with the heterologous PilT_Vc_^WA^. The relevant genotypes of each strain are indicated below each bar, which includes the status of the native *pilT/pilU* genes as well as the ectopically expressed retraction motor ATPases. The color of the bar corresponds to the species from which the ectopically expressed PilT motor ATPase was derived (green bars = *Vc* motors, orange bar = *Ps* motors). Data are from at least 3 independent biological replicates and shown as the mean ± SD.

**Fig. S3.**
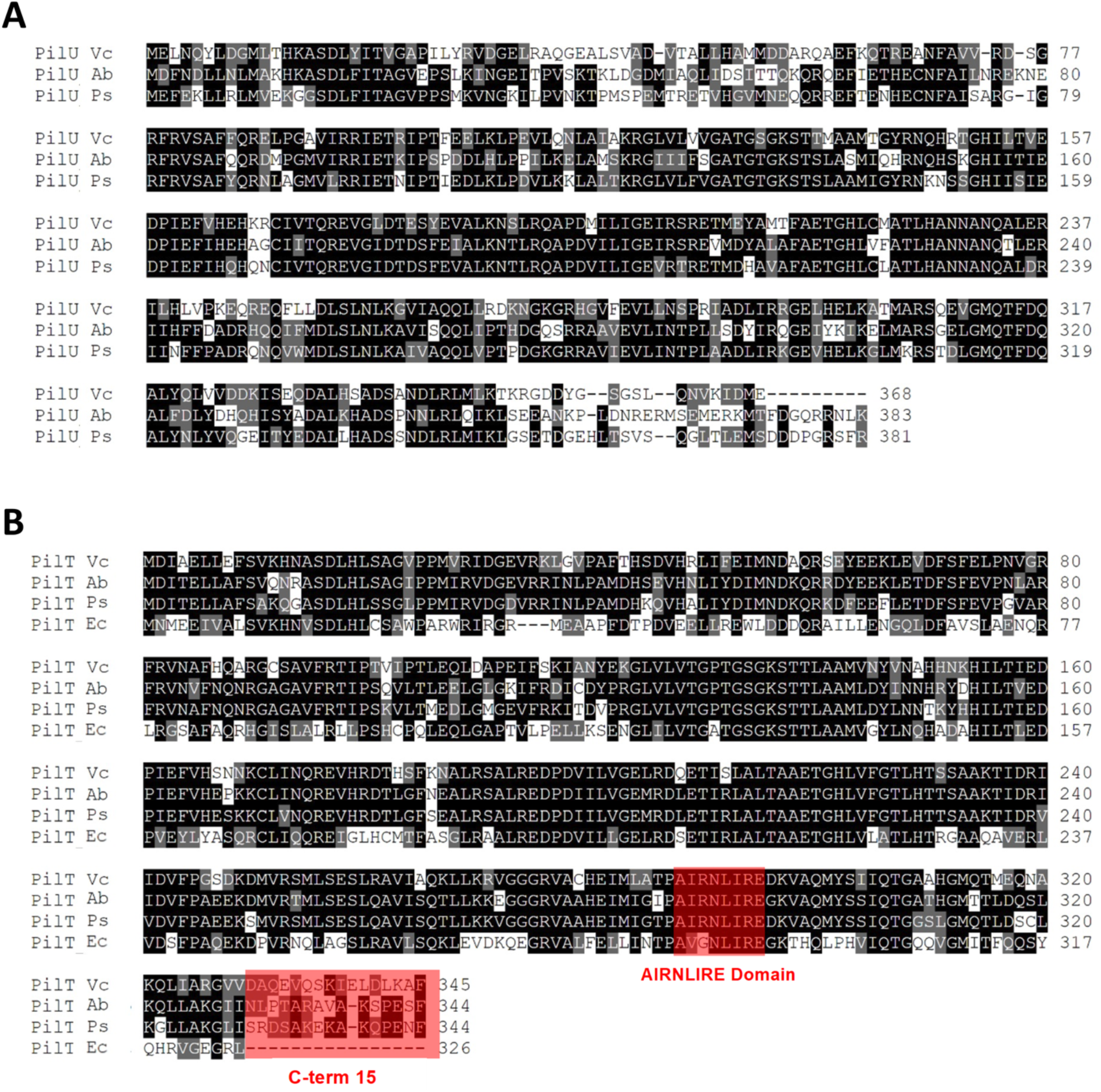
Alignment of PilU and PilT sequences from the four T4aP studied. Alignments of (**A**) PilU orthologs and (**B**) PilT orthologs. Residues highlighted in black are identical, while those highlighted in gray are similar. The AIRNLIRE and C-terminal 15 residue domains experimentally tested in this study are boxed in red. The GenBank Accession numbers for the protein sequences used are as follows: Vc, *Vibrio cholerae* PilT (AAF93635); Ab, *Acinetobacter baylyi* PilT (CAG67808); Ps, *Pseudomonas stutzeri* PilT (AFN76400); Ec, *Escherichia coli* PilT (NP_417425); Vc, *V. cholerae* PilU (AAF93636); Ab, *A. baylyi* PilU (CAG67807); Ps, *P. stutzeri* PilU (AFN76401).

**Fig. S4.**
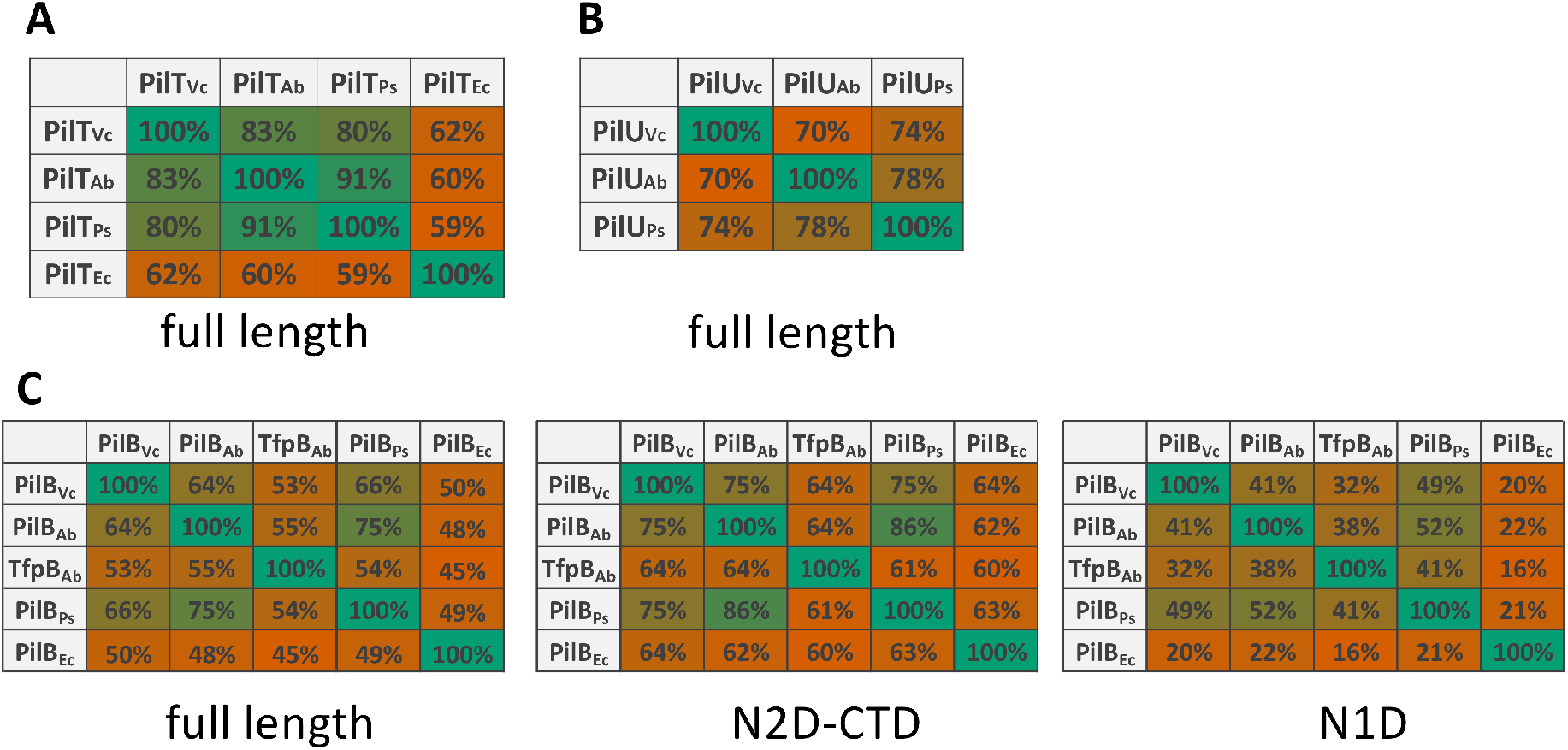
Pairwise comparisons of the indicated motor ATPases. Percent similarities are reported for pairwise comparisons of the indicated (**A**) PilT, (**B**) PilU, and (**C**) PilB/TfpB orthologs. The percent similarity is also denoted by the fill color of the box, where green represents high percent similarity and red represents lower percent similarity. PilB/TfpB orthologs were further analyzed for pairwise comparisons of the C-terminal domains that are shared between by extension and retraction motor ATPases (N2D and CTD) and the N-terminal domain (N1D) that is unique to extension motor ATPases (see **Fig. S5** for details on what portion of PilB represents the N1D vs N2D-CTD). The GenBank Accession numbers for the protein sequences used in these comparisons are as follows: Vc, *Vibrio cholerae* PilT (AAF93635); Ab, *Acinetobacter baylyi* PilT (CAG67808); Ps, *Pseudomonas stutzeri* PilT (AFN76400); Ec, *Escherichia coli* PilT (NP_417425); Vc, *V. cholerae* PilU (AAF93636); Ab, *A. baylyi* PilU (CAG67807); Ps, *P. stutzeri* PilU (AFN76401); Vc, *V. cholerae* PilB (AAF95567); Ab, *A. baylyi* PilB (CAG67312); Ab, *A. baylyi* TfpB (CAG69034); Ps, *P. stutzeri* PilB (AFN76867); Ec, *E. coli* PilB (AAC73218).

**Fig. S5.**
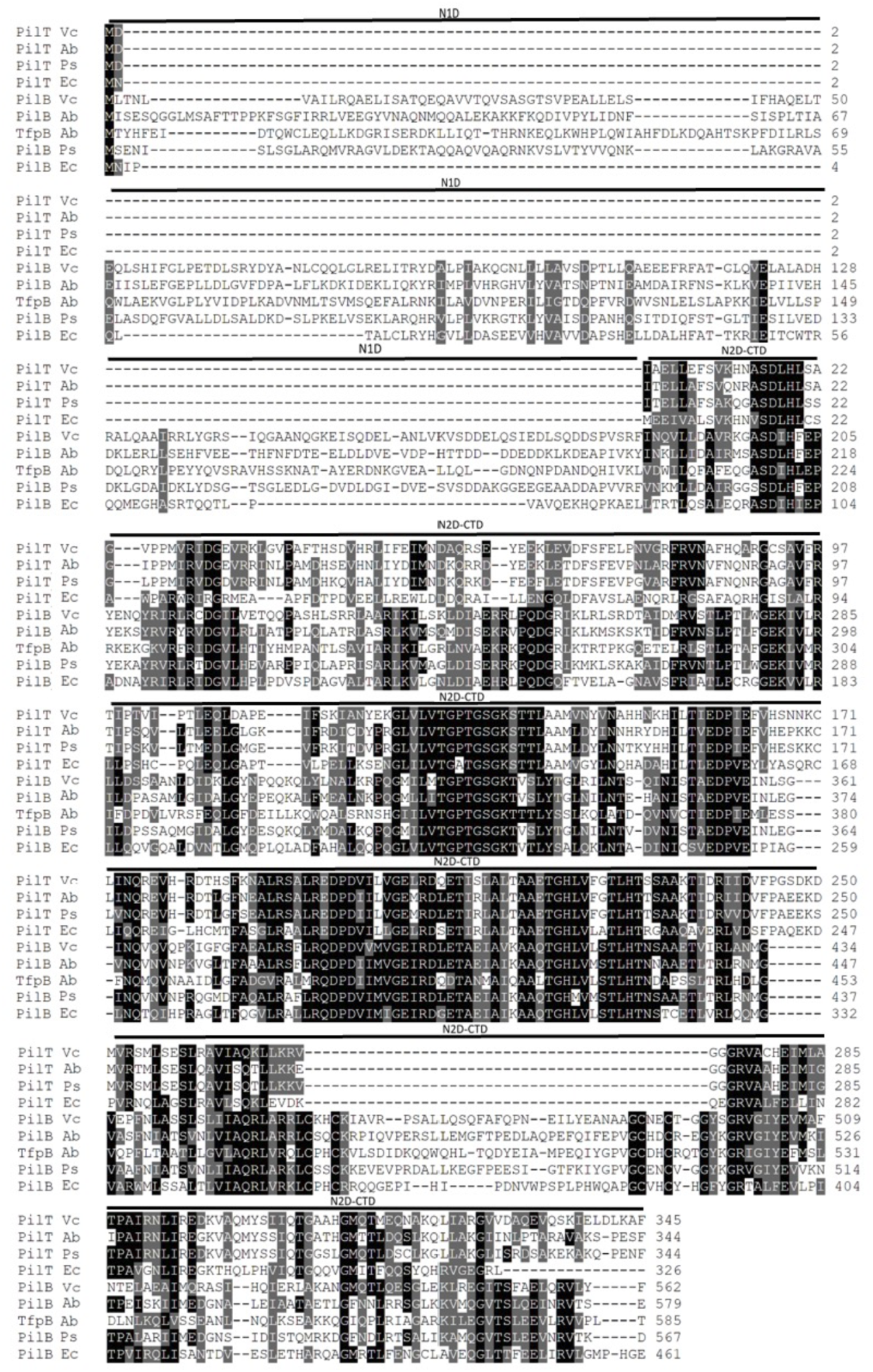
Alignment of the PilB/TfpB and PilT sequences from the T4aP studied. Alignments of PilB/TfpB orthologs and PilT orthologs. Residues highlighted in black are identical, while those highlighted in gray are similar. The unique N-terminal domain in PilB orthologs is denoted by the “N1D” bar above the sequence, while the two C-terminal domains shared by all motor ATPases are denoted by “N2D-CTD”. The GenBank Accession numbers for the protein sequences used are as follows: Vc, *Vibrio cholerae* PilT (AAF93635); Ab, *Acinetobacter baylyi* PilT (CAG67808); Ps, *Pseudomonas stutzeri* PilT (AFN76400); Ec, *Escherichia coli* PilT (NP_417425); Vc, *V. cholerae* PilB (AAF95567); Ab, *A. baylyi* PilB (CAG67312); Ab, *A. baylyi* TfpB (CAG69034); Ps, *P. stutzeri* PilB (AFN76867); Ec, *E. coli* PilB (AAC73218).

**Table S1.**
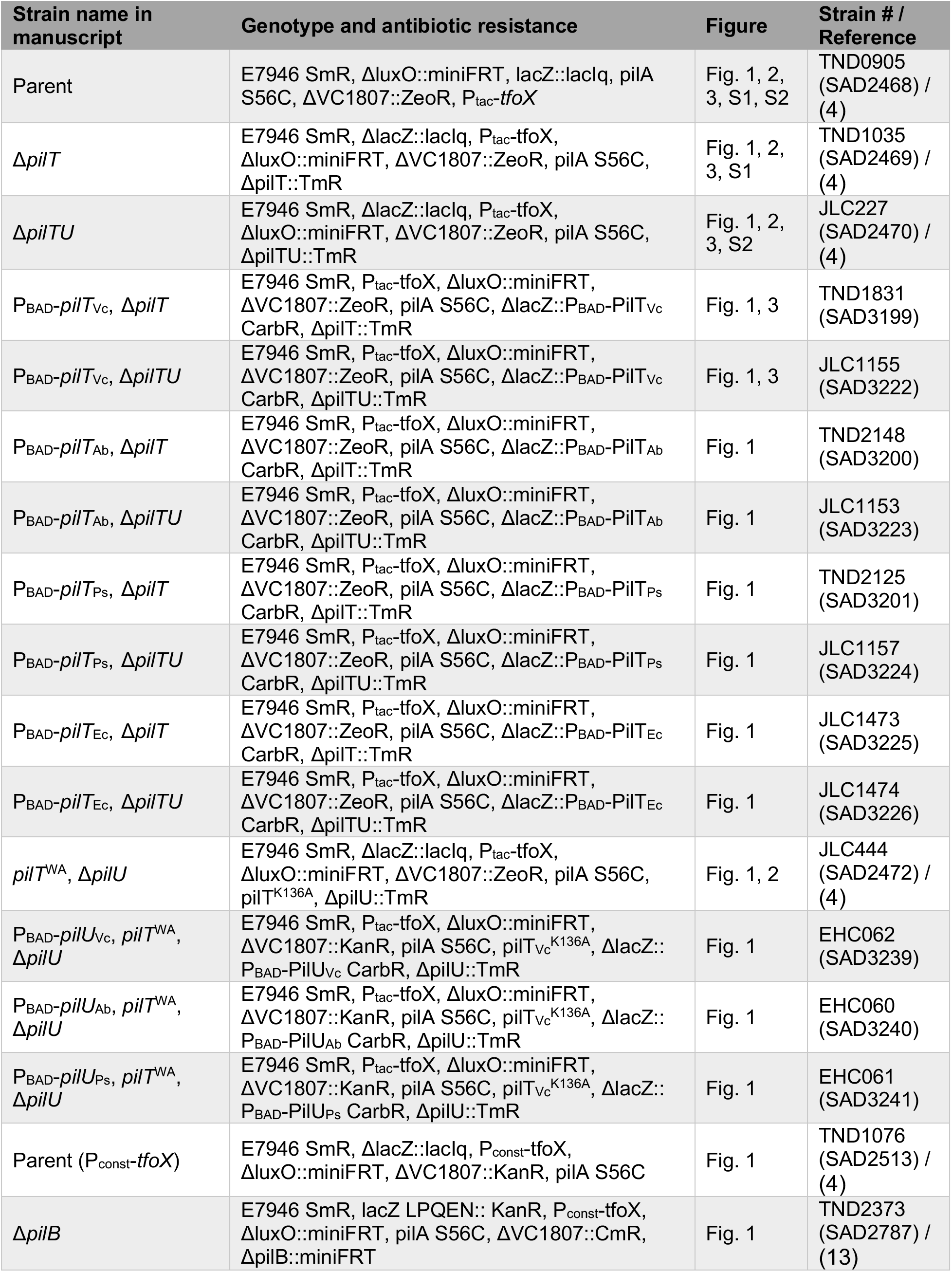
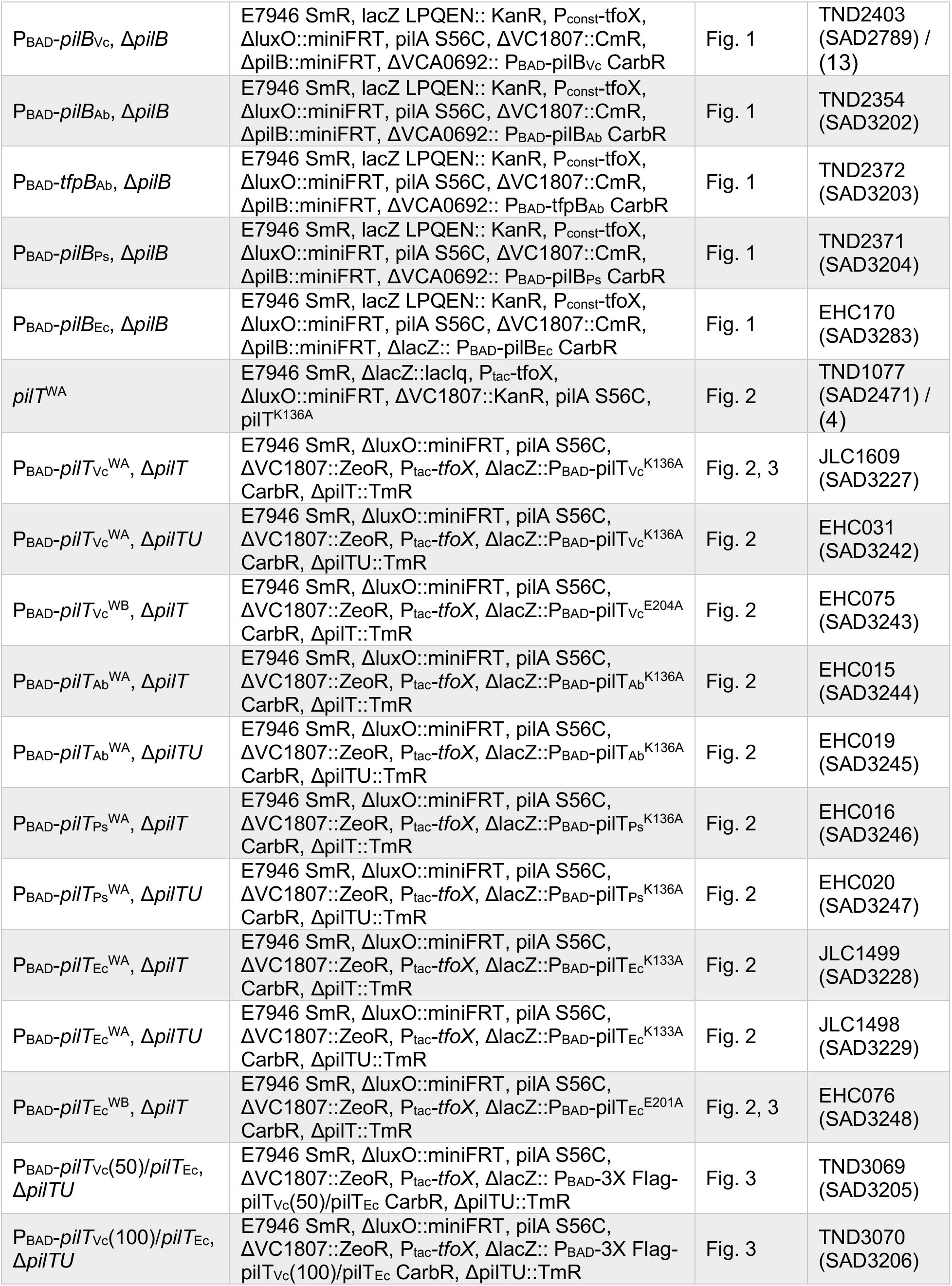
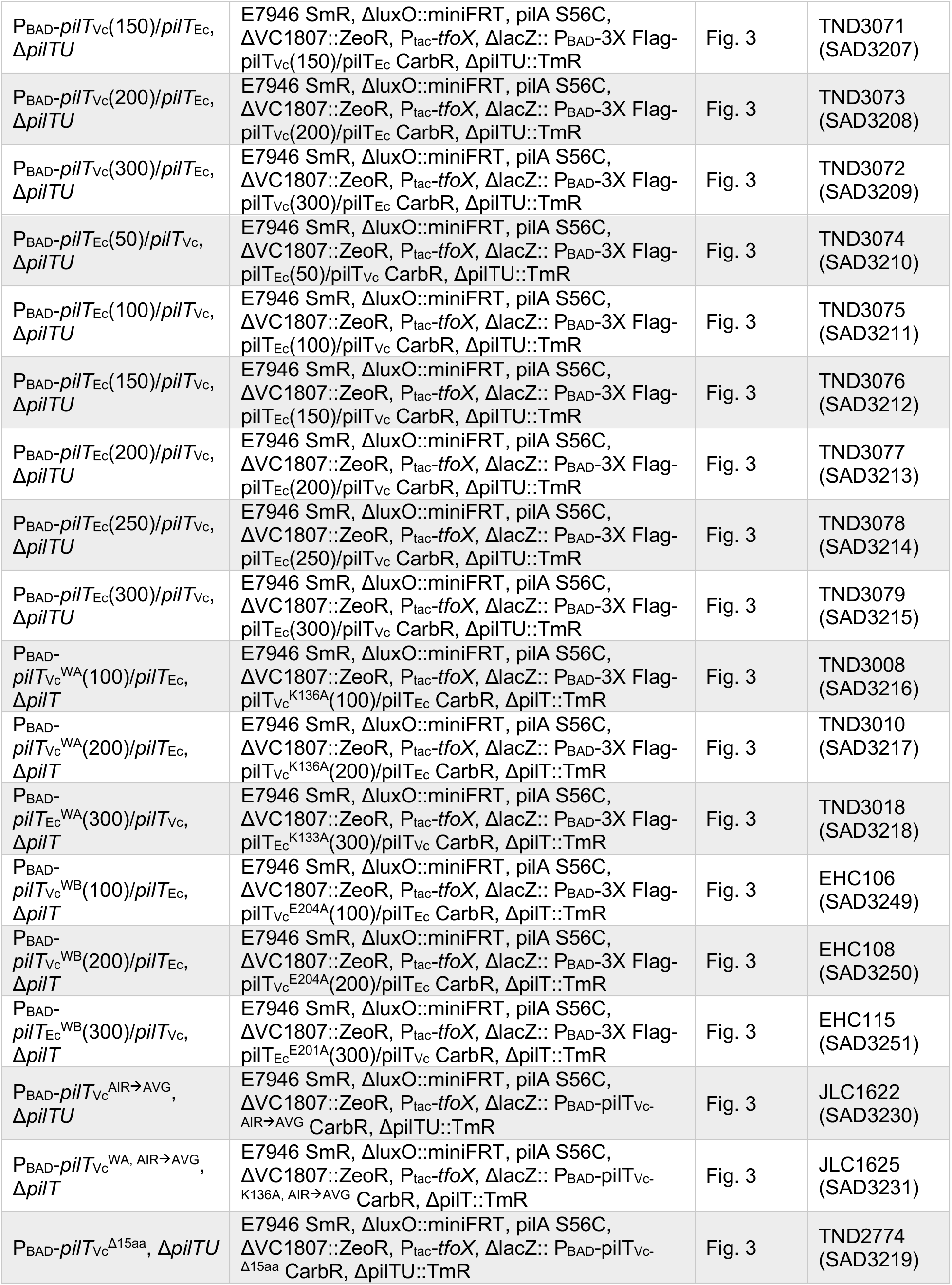
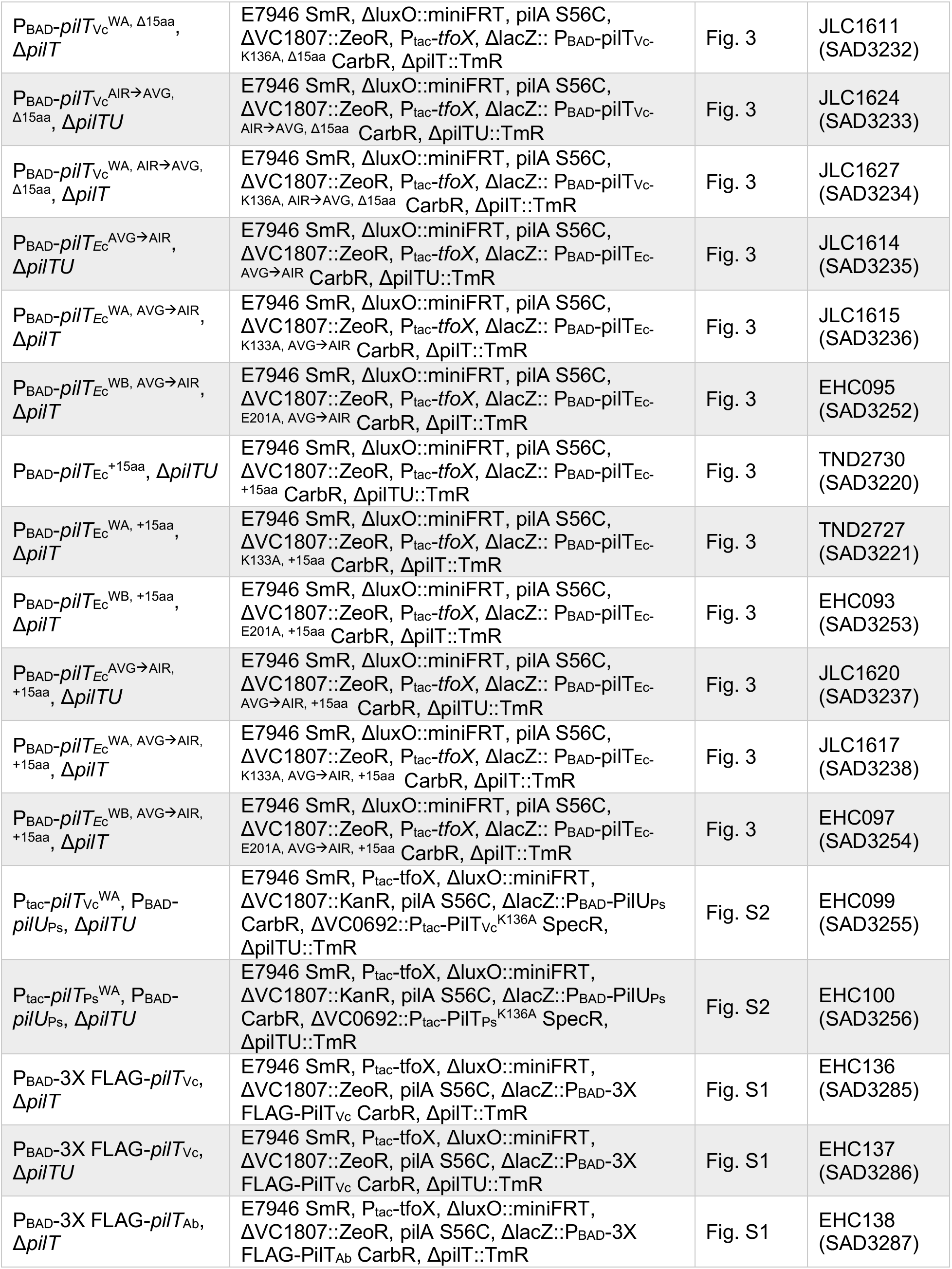
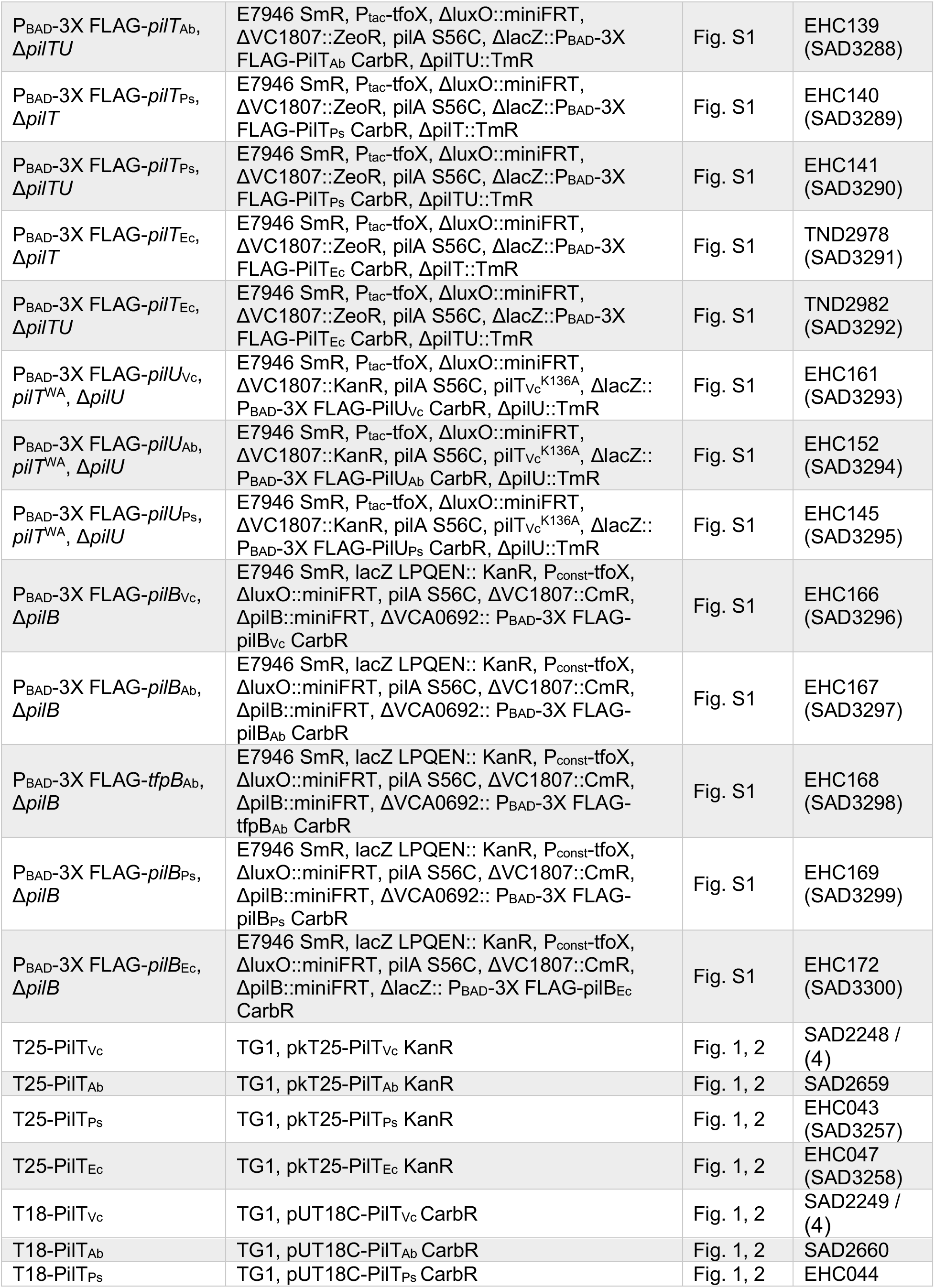
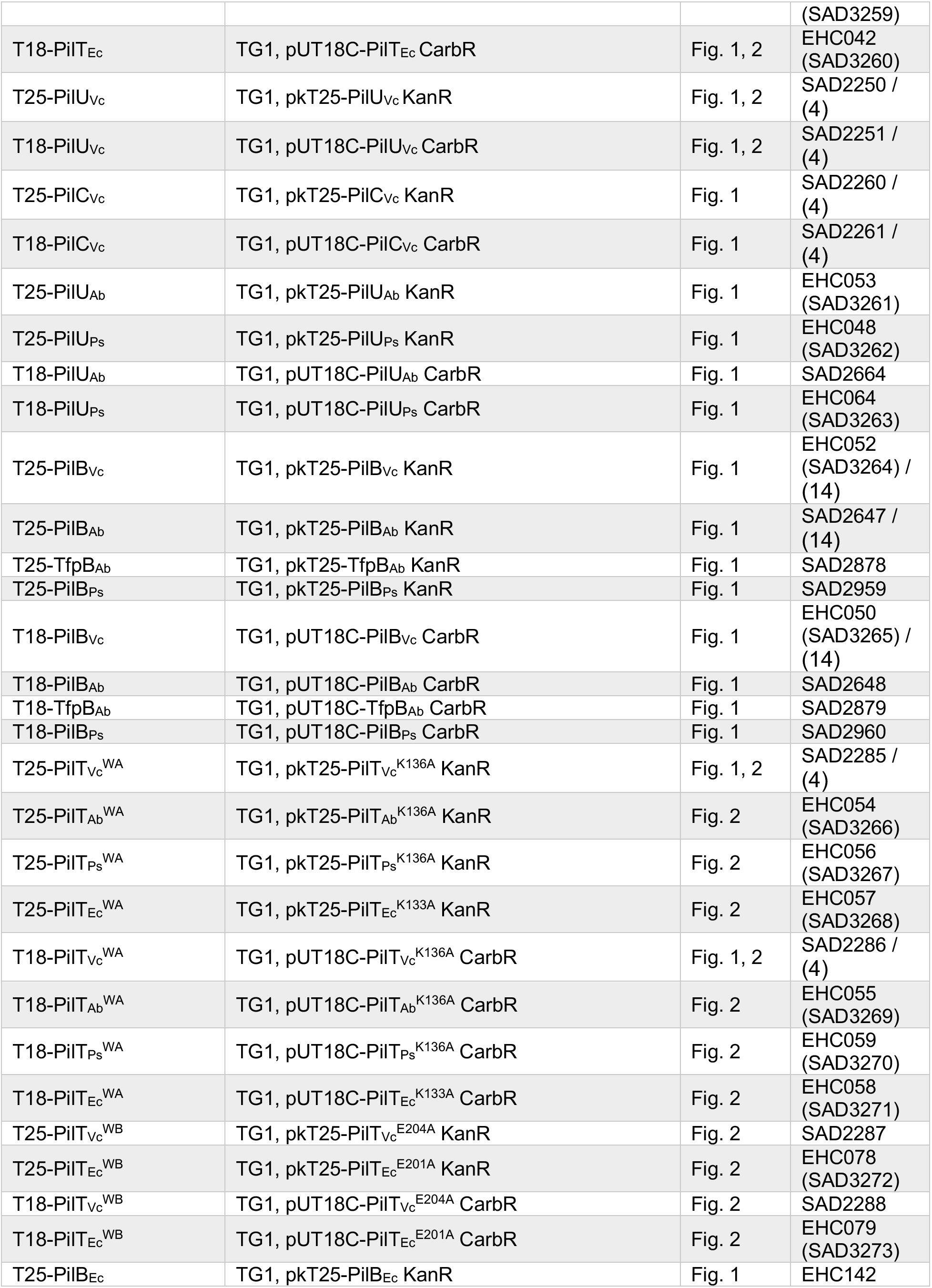
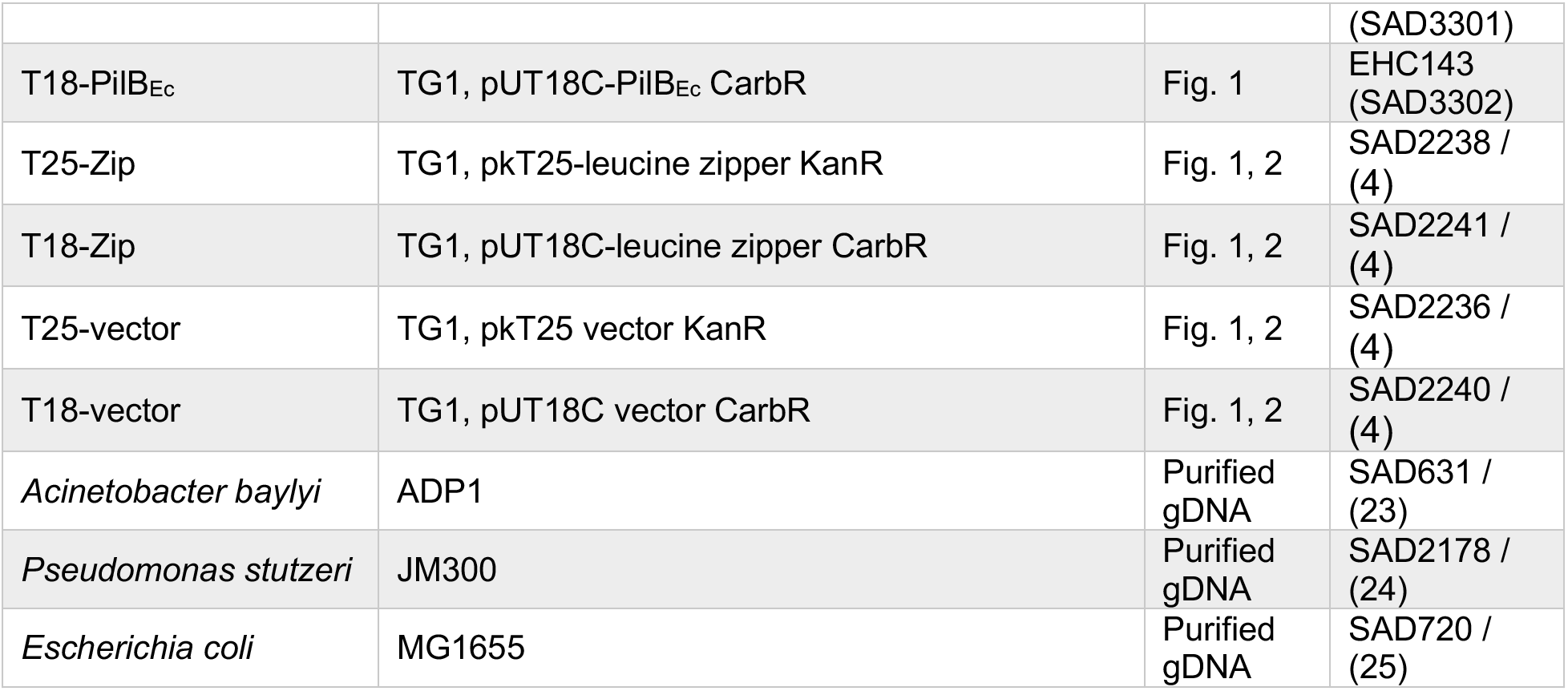
Strains used in this study.

**Table S2.**
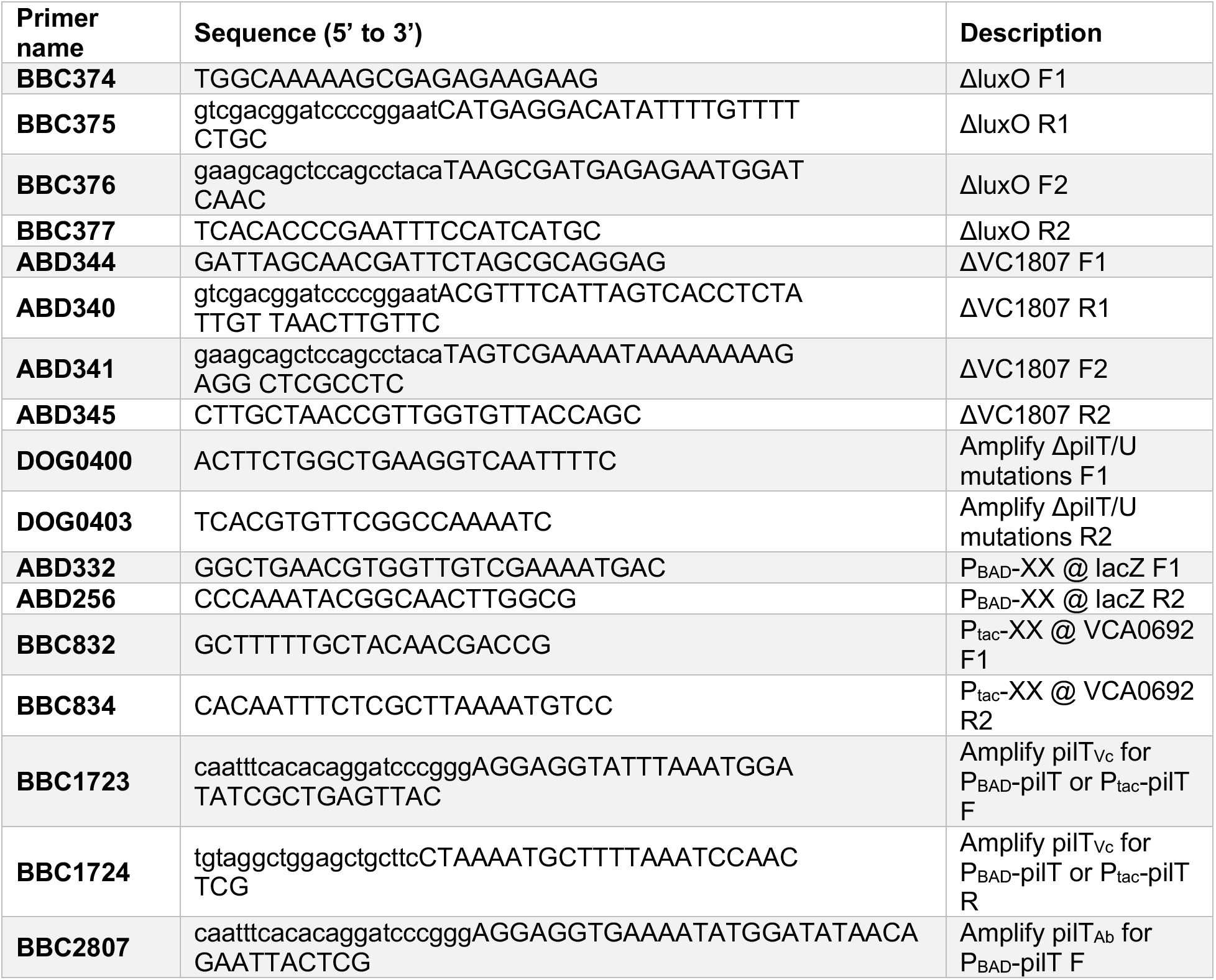
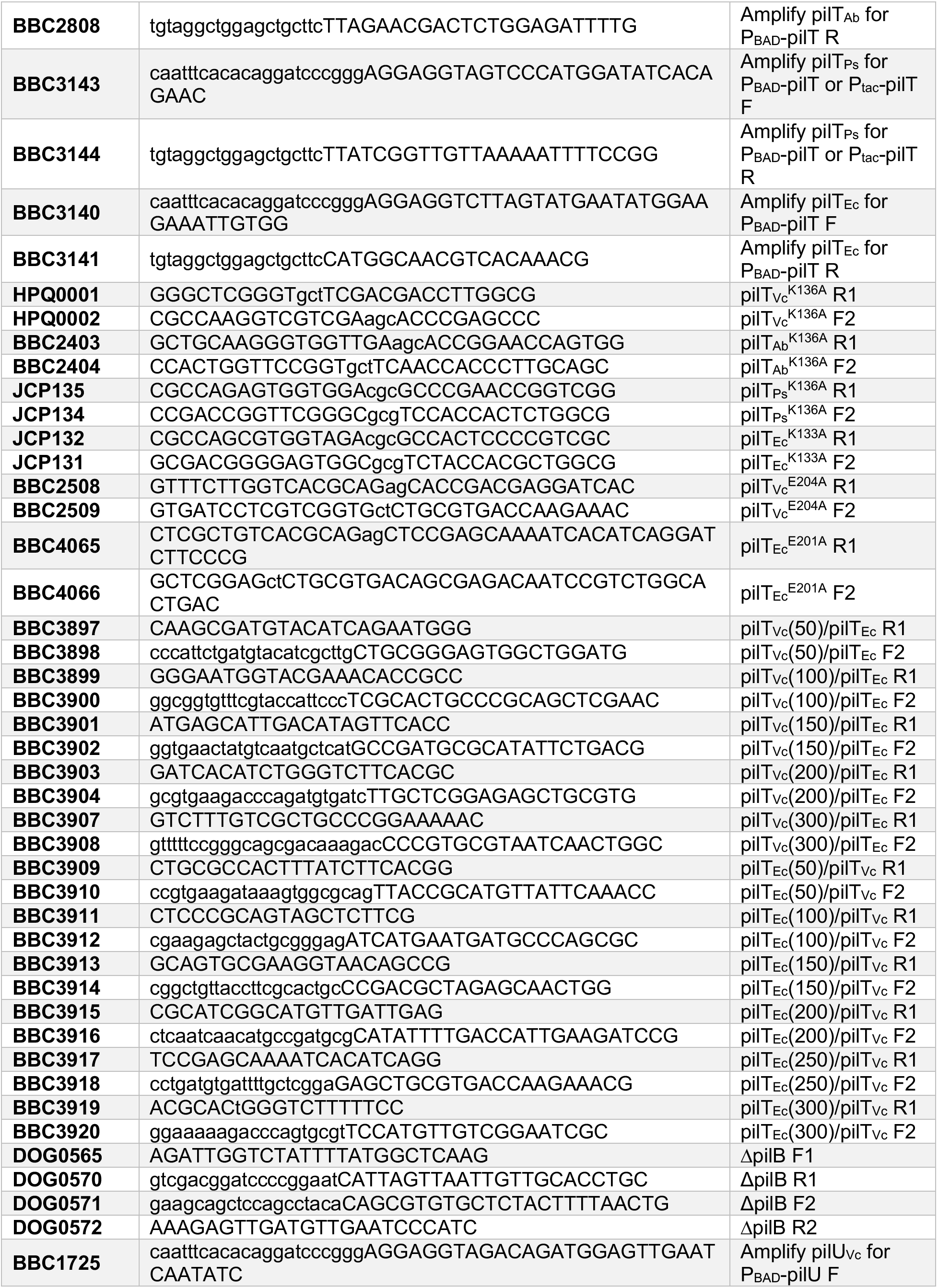
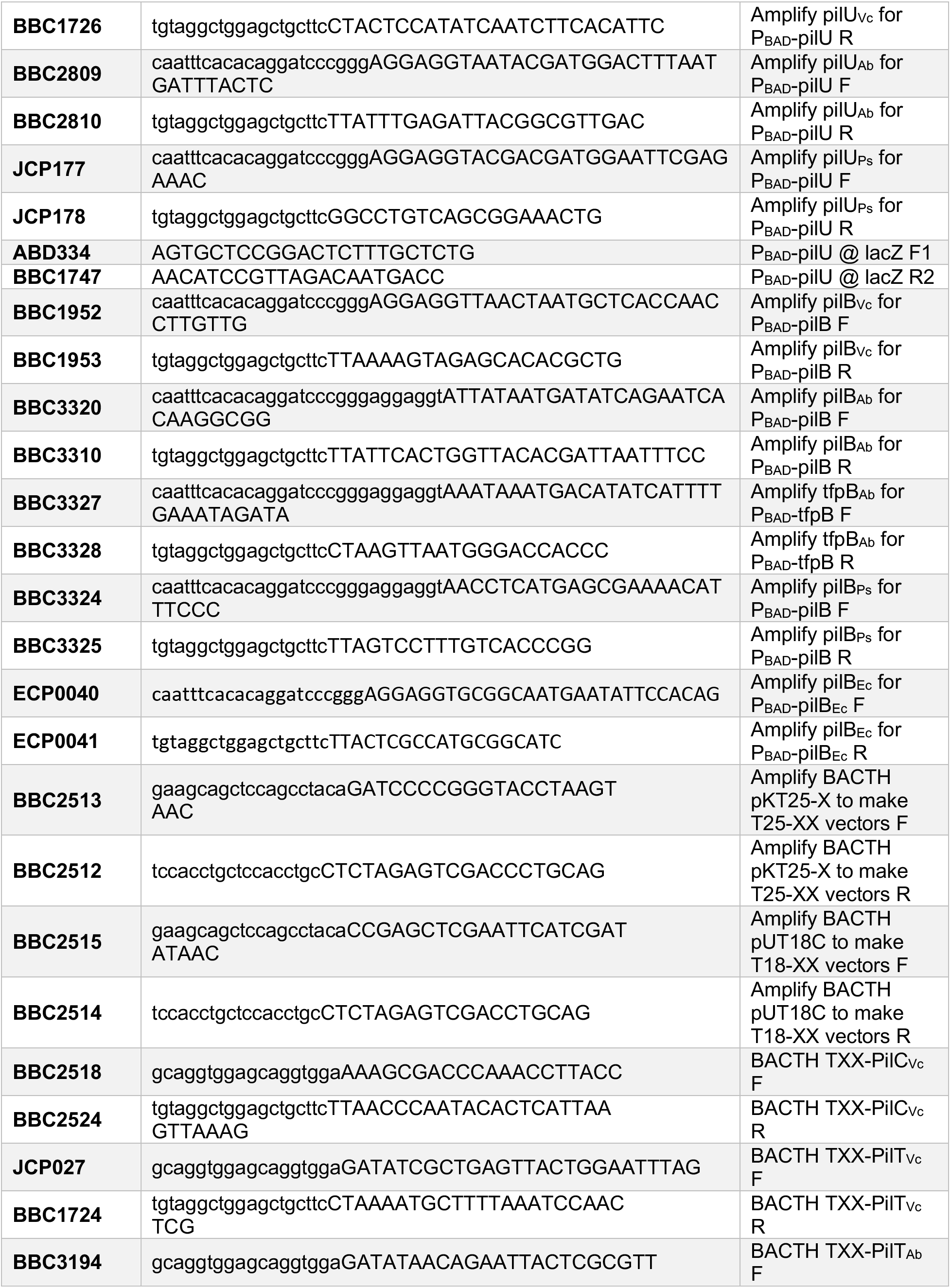
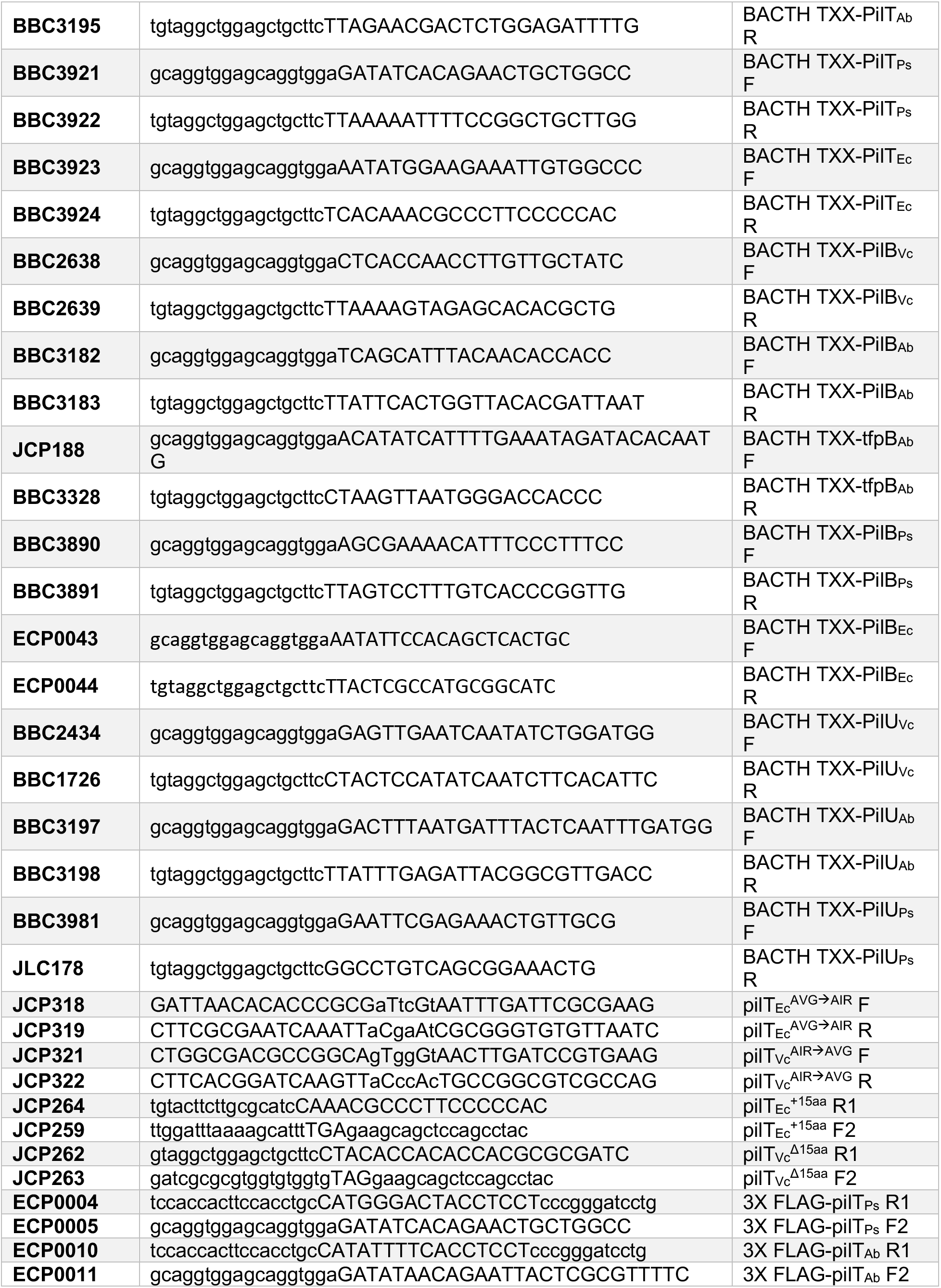
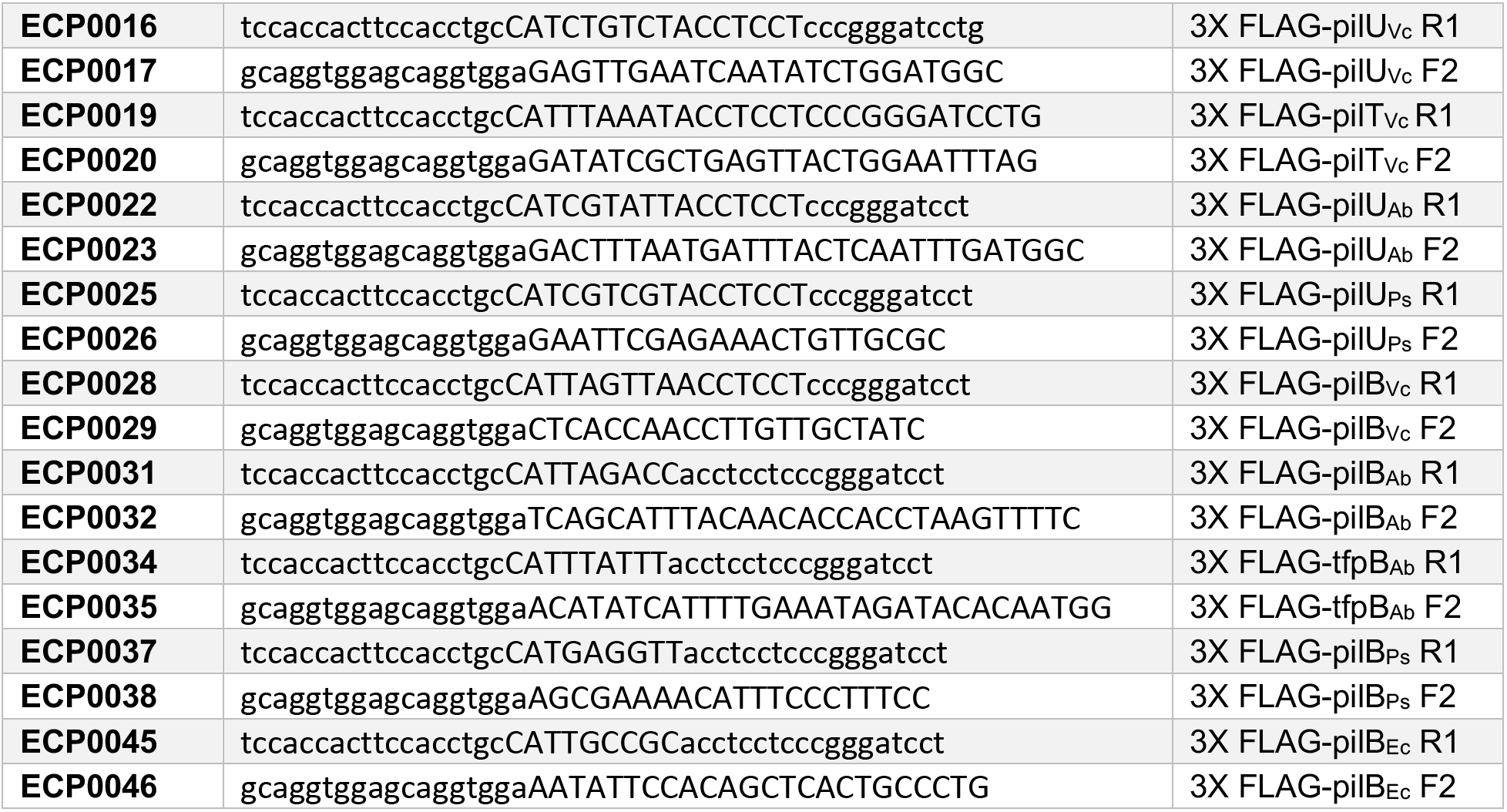
Primers used in this study.

